# Diagnosing linearity along the carbon cascade in terrestrial biosphere models

**DOI:** 10.1101/2025.09.01.673458

**Authors:** Huanyuan Zhang-Zheng, Vivek K. Arora, Peter Anthoni, Thomas A. M. Pugh, Atul K. Jain, Wenping Yuan, Yadvinder Malhi, Julia Nabel, Daniel S. Goll, Julia Pongratz, Benjamin Poulter, Anthony P. Walker, Sönke Zaehle, Jürgen Knauer, Etsushi Kato, Ruijie Ding, Minxue Tang, Stephen Sitch, Michael O’Sullivan, César Terrer, Hanqin Tian, Naiqing Pan, Pierre Friedlingstein, Akihiko Ito, Qing Sun, Jeanne Decayeux, Benjamin D. Stocker

## Abstract

Elevated carbon dioxide (eCO_2_) fertilises photosynthesis, driving an increase in terrestrial gross primary production (GPP). However, it is unclear how effectively increased GPP propagates along the “carbon (C) cascade” to increase net primary production (NPP) and vegetation C stocks (*C*_veg_) in different plant compartments. Vegetation models simulate divergent C cycle projections and have been criticised for being overly photosynthesis-driven (source-driven), neglecting processes that lead to non-linear behaviour in response to the GPP increase, which may attenuate (or amplify) changes in NPP and vegetation C stocks. Here, we introduce an analytical framework to diagnose linearity (*L*) of the land C cycle as the ratio of relative changes in linked fluxes and pools and apply it to outputs from 16 models of the TRENDY v11 ensemble. We found widely varying patterns in *L* across models and for the different links. Six models showed a clear dominance of larger relative changes in NPP than in GPP in global simulations (*L*_NPP:GPP_ >1 for >60% of gridcells), indicating increased carbon use efficiency under eCO_2_. Only three models had *L*_NPP:GPP_ < 1 for >60% of gridcells. Four models showed a clear dominance of larger relative changes in steady-state *C*_veg_ than in NPP, while five models showed an opposite pattern - in both cases with a large spread of *L*_Cveg*:NPP_ across gridcells within models. Three models showed a larger relative increase in root C than in *C*_veg_, while two models showed a clear dominance of the opposite pattern. Widely differing distributions of *L* across models and links reveal a strong influence of alternative process representations (nonlinear behaviour) in individual models. However, for all links, *L* deviations from 1 were roughly balanced across the model ensemble, leading to an overall linear behaviour of terrestrial C cycle representations.

## Introduction

Rising atmospheric CO_2_ has been stimulating photosynthesis of most land plants and has driven an increase in terrestrial gross primary production (GPP) by 13.5 ± 3.5% between 1981 and 2020 ^1^. Over the same time, the land biosphere has been a net carbon sink of 2.75 GtC yr^-1^ on average ^2–4^, varying annually from 0.36 to 4.1 GtC yr^-1^. This C sink partly offsets anthropogenic CO_2_ emissions and acts as a negative feedback against atmospheric CO_2_ growth, thereby slowing the pace of global warming ^5,6^. Although all organic carbon (C) stored in terrestrial systems ultimately originates from photosynthesis, it remains debated to what extent increases in GPP lead to increases in the terrestrial carbon sink ^3,7,8^. While the response of photosynthesis to CO_2_ plays out at the leaf level, the land C sink is most clearly inferred via the global C budget at the planetary scale ^9,10^. Establishing a causal link between the rise in GPP and the sustained positive land C sink is challenging because a large range of processes and feedbacks affect the link, and numerous fluxes and pools are involved in the propagation of GPP changes to changes in terrestrial C storage ^8,11^ (Box 1).

Each Dynamic Global Vegetation Model (DGVM) proposes its own way of representing this causal link. They resolve a variety of ecosystem processes from leaf to ecosystem scales and simulate their responses and feedbacks to rising CO_2_ and climate change ^12^. The representation of individual processes differs among DGVMs ^13–16^, leading to divergent simulations of C cycle changes during the past decades ^17^. However, common to all DGVMs is that they conceive the land C cycle dynamics as a *cascade* of C, representing that all C that cycles in terrestrial ecosystems is fuelled by photosynthesis and that a change in photosynthesis has cascading effects on all “downstream” C pools and fluxes between them ^18,19^. Following the notion of the C cascade, an increase in GPP is propagated to changes in net primary production (NPP), vegetation biomass, and soil carbon ^8,12,19^. If the land C cycle behaved as a linear system, any relative change in GPP would propagate to the same relative change in all downstream fluxes and pools when a new steady-state is hypothetically attained and in the absence of changes in external forcings (e.g., climate change) (Box 1). However, the linearity of this propagation is contentious, and it is an open question how effectively an increase in GPP, driven by elevated CO_2_, scales downstream fluxes and pools.

The central role of photosynthesis as the dominant driver of ecosystem C balance changes has been questioned, and it has been argued that such a “source-driven” conception of C cycle dynamics conflicts with key observations at different scales ^7,11,20,21^. For example, a decrease in *V*_cmax_, and leaf respiration under elevated CO₂ (eCO_2_) ^17,22–24^ could imply an increase in carbon use efficiency (CUE = NPP/GPP) and *L*_NPP:GPP_ > 1 ^17,22–24^. The link between NPP and biomass C storage could be affected by the tree growth-longevity trade-off ^25–30^, whereby accelerated tree growth (increased NPP) leads to an accelerated tree lifecycle (decreased vegetation turnover time τ) through amplified light competition, exclusion of short-statured plants from the canopy, and increased tree mortality in closed forest stands (*i.e.* more self-thinning) ^31^. These relationships have been identified from various tree and forest stand-level observations across large environmental gradients (tree rings, forest inventories), but it remains unclear whether similar relationships hold for the response to eCO₂ ^31^. If it does, the tree growth-longevity trade-off would imply decreased τ and *L*_Cveg*:NPP_ < 1 (Box 1). Furthermore, eCO₂ could lead to a shortage of soil nutrients, required for fixing C into biomass ^32–34^. Nutrient limitation could thus drive enhanced root respiration and/or labile C export to root symbionts ^24^. A shift of plant C allocation towards more short-lived fine root biomass and C exuded into the rhizosphere at the expense of allocation to long-lived aboveground woody biomass is often (but not always ^35^) recorded in ecosystem CO_2_ experiments ^17,22,24,36–41^. This suggests *L*_Croot:Cveg*_ > 1. An allocation shift towards more short-lived biomass under rising CO_2_ would imply that the total ecosystem-level biomass C stock increase is smaller compared to a case where the allocation remains unchanged (*L*_Cveg*:NPP_ < 1).

In recent years, DGVMs have been further developed to resolve several of these mechanisms. Additional processes have been implemented in the current generation of DGVMs. For example, among the 16 models used for annual updates of the global carbon budget ^2^, 10 models account for the effects of soil nitrogen on carbon cycling – in contrast to the first generation of DGVMs, where these fundamental resource limitations were not considered. Some of the N and P-resolving models simulate less NPP and shifts towards more belowground allocation after applying nutrient limitation ^42,43^. Three models resolve size-structured forest stand dynamics and vegetation demography as opposed to treating exclusively average-sized individuals as done in earlier DGVM generations and thus resolve some of the processes that lead to the growth rate-longevity trade-off ^44,45^. These additional process representations are expected to influence GPP propagation that fuels additional ecosystem C storage to some extent. Theoretically, the model dynamics should follow the model structure and process representations. However, the links between model structure and the effectively simulated dynamics of the C cascade are not straightforward to anticipate from the models manual due to feedback, non-linearity, threshold-type behaviour, and the ensuing state-dependency of effective functional relationships. Hence, it is unclear to what extent current generation DGVMs (still) simulate predominantly source-driven C cycle dynamics and what links along the C cascade – from GPP to NPP to vegetation carbon in different plant compartments – are responsible for a departure from linear systems dynamics.

Here, we introduce an analytical framework for diagnosing the ecosystem C dynamics from DGVMs and measuring the degree of linearity in the response of vegetation pools and fluxes to rising CO_2_ (Box 1). We focus on the links between GPP and NPP, between NPP and vegetation C, and between C in different plant compartments and total vegetation C, and quantify the relative responses and linearity terms of each variable and link from simulations of a set of current-generation DGVMs (TRENDY v11, S1). The further propagation to soil C storage has been investigated by a previous study ^41^ and thus is not analysed here. We quantitatively measure whether individual DGVMs behave in a linear manner (*L*=1, see Box 1). Considering that C-N coupled models and vegetation-demography models may behave differently from first-generation C-only and average-individual models, we also categorise models into distinct groups and summarise their respective behaviours. The framework applied here thus provides an approach for revealing the fundamental dynamics of DGVMs’ representation of the land C cycle and facilitates the comparison of model simulations and confronting them with findings from experiments and field observations.

### Box 1: The C cascade in terrestrial ecosystems - a linear system?

C enters ecosystems through photosynthesis, measured at the ecosystem level by GPP. Once in the system, it fuels a cascade of fluxes and pools. In a linear system, a constant fraction of C in a pool is transferred downstream, along the C cascade, per unit time, and the relative partitioning into multiple downstream pools remains constant. Such a representation of C dynamics in terrestrial ecosystems can be described by a linear system of equations ^19,46–48^. A linear representation of the C cascade implies that, given constant input fluxes, the hypothetical steady-state pool sizes are determined by their input flux; the time until the steady-state is approached scales with the turnover time of the respective pool; and the magnitudes of all fluxes scale linearly with the ultimate C source in the system – GPP.

To diagnose linearity (or deviations from linear behaviour) of DGVMs, we focus on three links in the C cascade. First, carbon use efficiency (CUE) represents the link between GPP and NPP (NPP = CUE × GPP). If CUE was constant, a relative change in GPP, here referred to as *R*_GPP_, induces an equal relative change in NPP (*R*_NPP_):

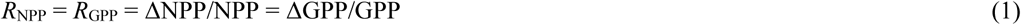

Where Δ denotes the absolute change of NPP and GPP, mostly due to eCO_2_ in this study. Second, a linear relationship between NPP and *C*_veg*_ follows from a first-order decay representation of the C dynamics, in which the loss of *C*_veg_ is proportional to the current value of *C*_veg_ and to the inverse of the turnover time (τ), while the gain of *C*_veg_ is given by NPP. The temporal change in *C*_veg_ given by the gain minus the loss, hence:

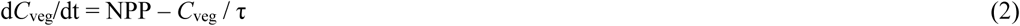

The steady-state vegetation C pool size (*C*_veg*_) is attained when dC_veg_/dt = 0 and is thus given by the product of NPP and the (effective) τ of C in that pool.

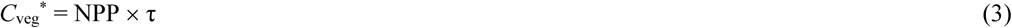

Hence, with τ being constant, a relative change in NPP induces an equal relative change in C_veg*_:

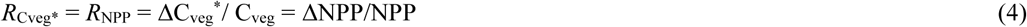

Note, that Eq. 4 describes relationships between NPP and *total* vegetation, representing the sum of C stored in individual plant compartments (leaves, wood, fine roots), and that τ represents an *effective* turnover rate, influenced by allocation to the different compartment. Since the typical turnover time of C varies strongly between plant compartments, the allocation (the relative partitioning) of NPP for building new biomass in different compartments is influential for the C dynamics and is implicitly assumed constant in Eq. 4.

Third, under constant allocation (and constant τ), the steady-state relative changes in root C and total vegetation C are identical.

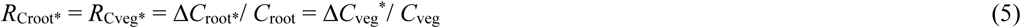

An analogous relationship applies for the other vegetation C compartments. That is, *C*_root_ may be replaced by *C*_leaf_ or *C*_wood_ in Eq. 5.

Linearity of these links can be measured by the ratio of relative changes and illustrated by regressing the relative change of the two variables of each link against each other. Departure of the ratio from 1 or points plotting off the 1:1 line in the regression indicates non-linear model behaviour. We thus consider the following three linearity metrics:

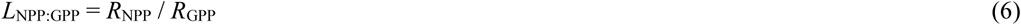

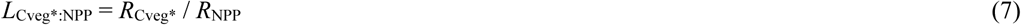

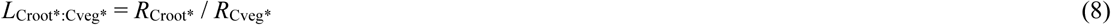

Environmental change is expected to affect CUE, τ, and allocation, e.g., through heat and drought-driven tree mortality, the effect of temperature on respiration rates, or the influence of reactive N deposition on growth and allocation. In contrast, CO_2_ exerts a direct control only on photosynthesis and GPP and has no direct influence on CUE, τ, and allocation. Thus, *R* and *L* terms can be diagnosed from model simulations where only CO2 is changing, while climate is held constant, to quantify the linearity of the terrestrial C cycle dynamics in response to changing CO_2_. As such, *L* = 1 indicates a perfectly linear behaviour. *L*_NPP:GPP_ = 1 indicates that CUE is not affected by eCO_2_. *L*_Cveg*:NPP_ = 1 implies constant vegetation turnover time and allocation patterns. *L*_Croot*: Cveg*_ = 1 indicates constant root:shoot ratios and above vs. belowground allocation patterns.

## Methods

### Model output processing

We obtained outputs of TRENDY v11 S1 simulations from 16 models ^2,49^. In these simulations, climate and land use were held constant at pre-industrial levels, while atmospheric CO_2_ concentrations changed as observed – from 285 ppm to 407 ppm – based on atmospheric and ice core measurements ^49^. In S1, nitrogen fertilization of agricultural land was fixed at pre-industrial levels, while varying nitrogen deposition on all land portions (agricultural and non-agricultural land) was prescribed from observations. Fire-enabled models allowed varying fire ignition probability. Using S1 simulations precludes climate-driven effects on ecosystem C dynamics (e.g. temperature change). TRENDY provides no simulations with varying CO_2_ and fixed N deposition, so this study includes both the effect of eCO_2_ and N deposition. Simulations were started from a dynamic steady-state in year 1700 and transiently covered the period up to the year 2022. We identified models as ‘C-N coupled models’ (n = 10) by referring to the supplementary material of this study ^50^. We identified models as ‘vegetation demography models’ (n = 3) by consulting modellers. We considered vegetation demography models as those that simulate tree-size dependent demographic rates (growth, mortality, fecundity) and distinguish trees (or cohorts of trees) of different sizes within each plant functional type (PFT) and within a forest stand. ‘Vegetation demography models’ include CABLE-POP, LPJ-GUESS and SDGVM. A list of ‘C-N coupled models’ could be found in the Supplementary Data.

Although land use and climate do not change in TRENDY S1 simulations, the land cover fraction of different PFTs may change in response to rising CO_2_ (and N deposition), or in response to changing wildfire in models that simulate dynamic PFT distributions. In particular, shifts between grasses and trees are accompanied by changes in aboveground biomass stocks even if NPP remains at similar levels. This is due to changes in the fraction of C allocated to (long-lived) wood that is zero for grasses. For models with dynamic PFT (LPJ-GUESS, JULES, and LPX-Bern), we thus excluded gridcells for which the land cover fraction of the primary PFT changed between simulation start and end.

Sparsely vegetated regions (e.g., arid deserts and tundra) may have extreme or no relative changes in carbon flux. We noticed that most of these low-biomass gridcells lead to *L*_Cveg*:NPP_ = 0 (Fig. S20). Excluding gridcells with *C*_veg_ below the 5% quantile could preclude many *L*_Cveg*:NPP_ = 0 (Fig. S20). Under very sparsely vegetated conditions, processes can be very different from those described in Box 1. It’s clearer to remove these pixels to facilitate interpretation. We thus excluded gridcells with *C*_veg_ below the 5% quantile for figures associated with *C*_veg*_ and *C*_veg_.

For *L*_Croot:Cveg_, LPJ-GUESS, DLEM, JSBACH and LPJ were excluded from the analysis because *C*_root_ of these models was not reported. For the same reason, some models were excluded from analysing *L*_Cleaf:Cveg*_ and *L*_Cwood:Cveg_. JULES and IBIS include coarse root biomass in *C*_wood_, while other models include it in *C*_root_ (Fig. S19). Hence, figures associated with *C*_root_ should be interpreted with this in mind.

### Quantifying linearity

For a given model, the relative change *R* of variable *X* was calculated for each gridcell *i* as:

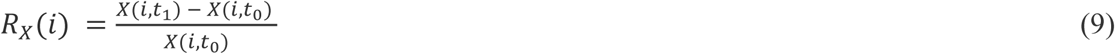

where *X*(*i*, *t*_0_) is the mean of *X* over the first 10 simulation years, 1700 to 1709 (*t*_0_) and *X*(*i*, *t*_1_) is the mean over years 2013 to 2022 (*t*_1_). As variables *X*, we considered GPP, NPP, total biomass C (*C*_veg_), root biomass C (*C*_root_) and other components of biomass. The linearity term *L* of the link between two variables *X* and *Y* was calculated as the ratio of their relative changes

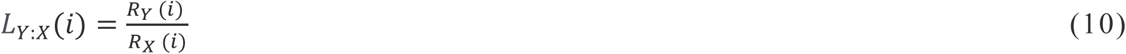

We quantified *L* for the link between GPP and NPP (*L*_NPP:GPP_), between NPP and *C*_veg_ (*L*_Cveg*:NPP_), and between *C*_veg_ and C in individual plant biomass compartments (for example *L*_Croot:Cveg_).

Linearity (*L*) of each DGVM should be calculated by quantifying the ratio of relative changes in connected pools and fluxes at their respective steady-state (Box 1). However, the model outputs analysed here are from transient simulations, forced by gradual CO_2_ changes that increased from a stable preindustrial level (285 ppm in 1700) in an accelerating manner to its current level (407 ppm in 2022). At this time scale, pools with turnover time scales on the order of decades and longer cannot be assumed to track their steady-state size during the simulations. For example, most of the vegetation carbon (*C*_veg_) is contained in woody biomass, which has a turnover time of decades to centuries ^29^. Thus, when dealing with links between fluxes and pools (e.g., *L*_Cveg*:NPP_), the steady-state of such pools needs to be estimated from the reported transient state.

By re-arranging Eq. 2 (Box 1), the turnover rate can be estimated from NPP, the current size and the temporal change of a C pool:

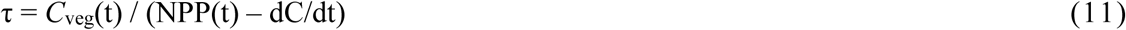

By combining Eq. 11 and Eq. 3, we get an expression for estimating of the steady-state vegetation C pool size from the transient dynamics:

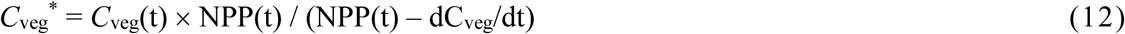

Eq. 12 was applied here for estimating the steady-state vegetation C pool size *C*_veg_* from the transient simulation outputs, whereby *C*_veg_(t) in Eq. 12 was taken as the mean of *C*_veg_ across years 2013 to 2022. dC_veg_/dt in Eq. 12 was taken as (*C*_veg_(2022) − *C*_veg_(2013))/10. NPP(t) was calculated as the mean across 2013 to 2022.

GPP is controlled by the physiology of photosynthesis, which has a relatively rapid response to CO_2_, and by the structure of the canopy. Leaf area changes in response to the environment evolve over seasonal time scales and respond to altered CO_2_ within years or faster, as seen for example at the Duke FACE experiment ^51^. We thus did not apply the steady-state correction (Eq. 12) on GPP and used unmodified model outputs for respective analyses. NPP is equal to GPP minus autotrophic respiration (*R*_a_). *R*_a_ contains growth respiration (which scales with NPP), fine root, leaf, and sapwood maintenance respiration. Fine root and leaf productivity and biomass in models respond on fast time scales (seasons to years), which should thus equilibrate correspondingly fast. Sapwood respiration may take longer to equilibrate if taken to be proportional to sapwood mass, if sapwood mass scales with total woody biomass, and given that woody biomass has a decadal to centennial response time scale. However, sapwood respiration is only a relatively small fraction of *R*_a_ ^52^. Based on the above, we neglected non-equilibrium effects for GPP and NPP. Similarly, for *C*_leaf_ (respective links shown in supplementary results), we assumed equilibrium given the short lifetime of leaf biomass (months to years) ^53,54^. Thus, *L*_Cleaf:Cveg*_ is presented.

For the link between *C*_root_ and *C*_veg_, and for the link between *C*_wood_ and *C*_veg_, the steady-state correction would be ideal, because these carbon pools have long turnover time, from several decades to 100 years (Xue *et al.*, 2017; Yu *et al.*, 2023). However, we could not apply the steady-state correction (Eq. 12) because DGVMs did not report root NPP and woody NPP. We chose to compare these to uncorrected *C*_veg_ because they both have decadal turnover time. The non-equilibrium effect may cancel out when regressing them for *L.* Therefore, we analysed *L*_Croot:Cveg_ and *L*_Cwood:Cveg_. We also provide results where we applied the steady-state correction on *C*_veg_ for an alternative evaluation of *L*_Croot:Cveg*_ (Fig. S10). We found very similar patterns to the uncorrected version (*L*_Croot:Cveg_) (Fig. 4), but with a general tendency for lower values.

### Illustration of linearity

The relative increase terms *R* and linearity terms *L* are evaluated for each link, gridcell, and model. We provide results at different levels of aggregation, including spatial maps, density scatter plots, and histograms of the distributions of *L* for each link and model and a summary table of median values for *L*.

For the spatial maps, we calculated the median of *L* terms from all models for each gridcell where the median represents overall *L* across models. As DGVMs vary in resolution, we first resampled model outputs to a coarser common resolution (1 degree in longitude and latitude). For the density scatter plot and histogram of the term *L*, such resampling was not suitable since resampling would reduce the range of values (extreme values would be smoothed out). The outputs of the model with the coarsest spatial resolution (VISIT) were provided for a total of 58,697 land gridcells. Therefore, we randomly sampled 58,697 gridcells from each model and pooled all values from all models to present the ‘overall’ pattern (e.g. Fig. 1a, 1b) and the ‘all’ panel in supplementary information. The random sampling thus ensures that outputs from low spatial resolution models are not underrepresented in the analysis of pooled data from multiple models.

**Figure 1.**
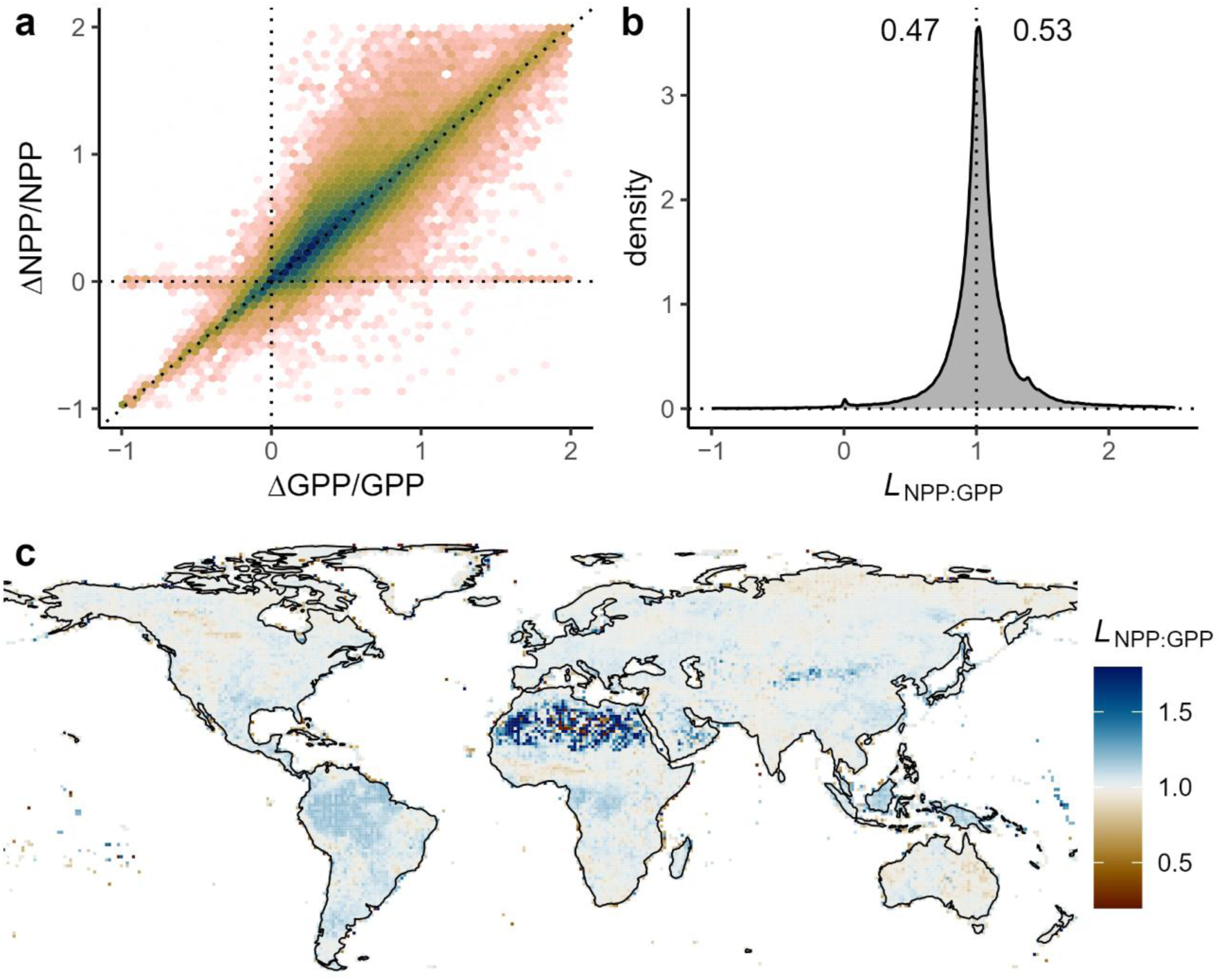
The relationship between the relative change of net primary productivity (*R*NPP = ΔNPP/NPP) and the relative change of gross primary productivity (*R*GPP = ΔGPP/GPP) under rising CO2 and changing N deposition, aggregated across models. (a) Density scatter plot of *R*NPP vs. *RG*PP values across all gridcell, pooled from balanced samples of all models. Dark colours denote high density, while bright colours denote low density. The dotted lines are the y=0 line and the y=x line. (b) Density of the distribution of *L*NPP:GPP (=*R*NPP/*R*GPP) values, pooled from balanced samples of all models. (c) Spatial pattern of *L*NPP:GPP, shown as the median per gridcell across models.

## Results

### The link between GPP and NPP

Across models and gridcells within models, there was a clear linear relationship between changes in GPP and NPP with a slight tendency towards larger relative increases in NPP than in GPP under rising CO_2_ (Fig. 1). The magnitudes of relative changes in GPP (*R*_GPP_) varied across models. Some models simulated relatively small increases (SDGVM: 13% [interquartile range across gridcells: 7%]; CLASSIC: 11% [IQR 23%]), while others simulated larger increases (e.g., ISBA-CTRIP: 39% [IQR 20%], ORCHIDEE: 31% [IQR 40%], ISAM: 31% [15%], and CABLE-POP: 31% [IQR 26%]) (Supplementary Data). However, while *R_G_*_PP_ and *R*_NPP_ varied substantially across models, their ratio (L) was tightly conserved. Pooled outputs showed narrow clustering of *R_G_*_PP_ - *R*_NPP_ along the 1:1 line (Fig. 1a) and a tight distribution of *L*_NPP:GPP_ around unity (1.02 [IQR: 0.22]; Fig. 2).

**Figure 2.**
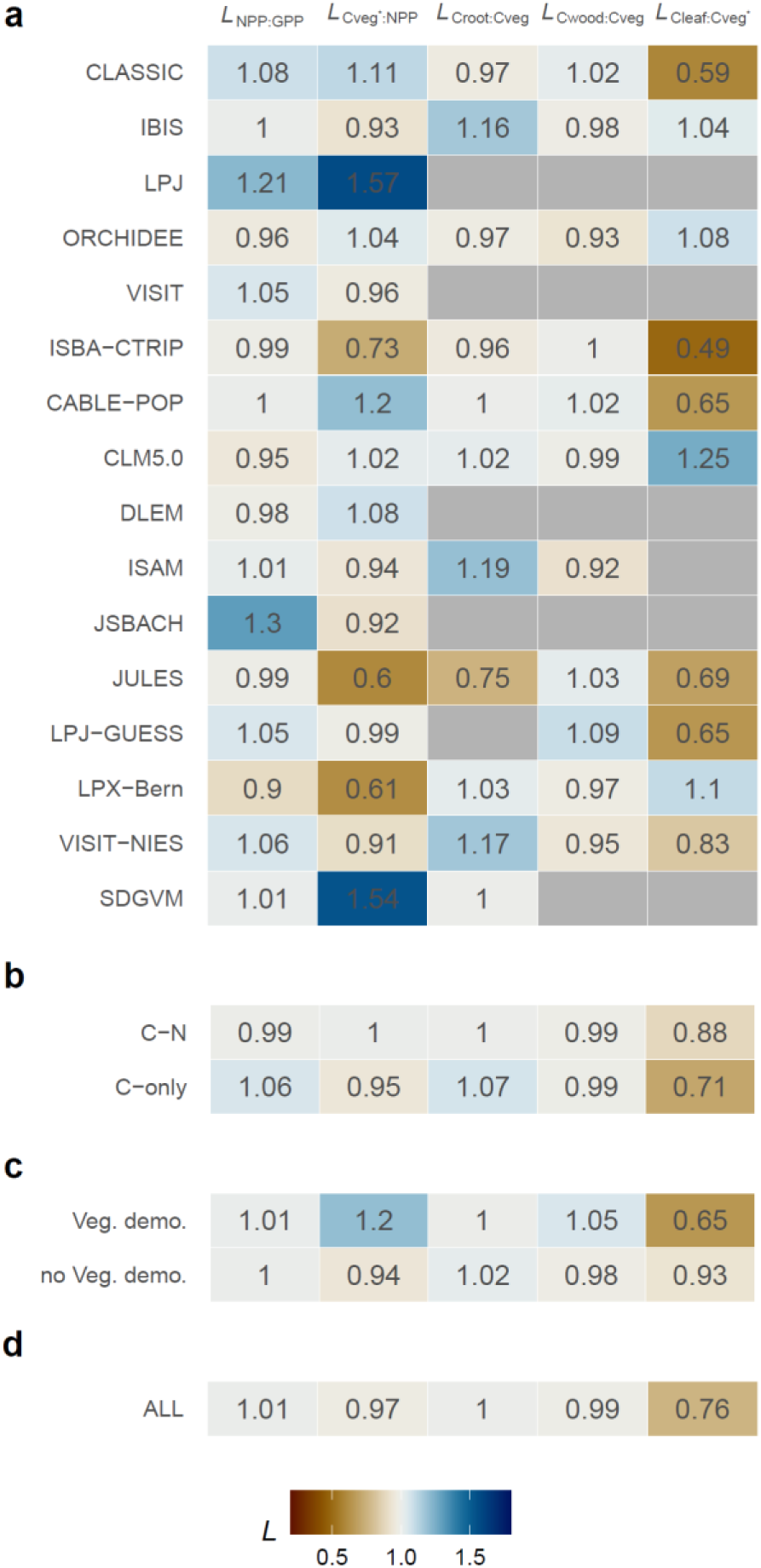
Overview of linearity terms *L*NPP:GPP, *L*Cveg*:NPP, and *L*Croot:Cveg across models. We show the median of all gridcells for each model (a). We also show linearity terms *L* as a median across all models (d) and model groups according to whether the model considered carbon nitrogen coupling (C-N) (b) and vegetation demography (Veg. Demo.) (c). Values of the linearity terms are given as annotations and accordingly coded by colours. The interquartile range (IQR) ancillary to the median is provided in Supplementary Data. Grey cells in (a) indicate missing values due to the non-availability of compartment-specific vegetation C outputs. See supplementary data for models’ C-N and Veg. Demo. grouping information.

Although the simulated link between GPP and NPP is predominantly linear (*L*_NPP:GPP_ 1) overall, individual models show strong deviations from this pattern (Figs. S1 and S2). Several models (CLASSIC, LPJ, LPJ-GUESS, SDGVM, VISIT, JSBACH) exhibit patterns of *L*_NPP:GPP_ > 1 for more than 50% of gridcells (Fig. S1). JSBACH shows the extreme where 76% of all gridcells exhibit *L*_NPP:GPP_ > 1. Several models show the opposite, including LPX-Bern, ORCHIDEE and DLEM. In LPX-Bern, only 25% exhibit *L*_NPP:GPP_ > 1 and a substantial portion of gridcells exhibit zero NPP change despite changes in GPP (Fig. S1, S2).

Geographical patterns emerge on average across all models, with a tendency towards *L*_NPP:GPP_ > 1 in many moist tropical regions and *L*_NPP:GPP_ < 1 in drier regions high-northern latitudes (Fig 1c). However, patterns vary strongly across models (Fig. S3). For example, JSBACH exhibits a clear tendency of *L*_NPP:GPP_ > 1 across the globe, but not for tropical rainforests. ORCHIDEE simulates an opposite pattern with a tendency for *L*_NPP:GPP_ < 1 across the globe, but > 1 in for tropical rainforests. In several models (IBIS, VISIT, VISIT-NIES), *L*_NPP:GPP_ is rather uniform across the globe or doesn’t exhibit a clear latitudinal pattern.

Across most models without C-N coupling, there is a tendency for *L*_NPP:GPP_ > 1 with median *L* = 1.06 (Fig. 2). Five out of the 10 C-N coupled models simulated *L*_NPP:GPP_ < 1, while the other five models simulated an opposite tendency, yielding median *L* = 0.99. We found no systematic differences between models incorporating vegetation demography (*L* = 1.01) versus average-individual models (*L* = 1.00).

### The link between NPP and vegetation carbon

The overall median of *L*_Cveg*:NPP_ is around unity at 0.97 [IQR 0.73] (Fig. 3). Unity indicates that, on average, the change in vegetation carbon tends to be linearly related to changes in NPP. However, the distribution of *L*_Cveg*:NPP_ was much wider than that of *L*_NPP:GPP_, owing to more spatial variation and more inter-model disagreement (Fig. 3 compared to Fig. 1). Although most individual models clearly deviate from an overall linear relationship between NPP and *C*_veg_, the general pattern of the results pooled from all models indicates a peak of the distribution of *L*_Cveg*:NPP_ at unity and a roughly balanced share of pixels across models indicating *L*_Cveg*:NPP_ > 1 vs. *L*_Cveg*:NPP_ < 1 (Fig. 3). The patterns in *R*_Cveg*_ versus *R*_NPP_ were very diverse (Fig. S4 and S5). For example, ISBA-CTRIP exhibited overall no clear relationship between the magnitudes of relative changes in NPP and *C*_veg_ and a clear dominance of *L*_Cveg*:NPP_ < 1. In ORCHIDEE, there was a tendency of *L*_Cveg*:NPP_>1 for small values of *R*_NPP_ and *L*_Cveg*:NPP_ < 1 for large values of *R*_NPP_. Several models had *L*_Cveg*:NPP_> 1 for a clear majority (>60%) of gridcells (CABLE-POP, DLEM, LPJ, SDGVM). Among them, DLEM, LPJ, and CLM5.0 had a substantial portion of gridcells for which *R*_Cveg*_ was much larger than *R*_NPP_ (Fig. S5). In contrast, several models have *L*_Cveg*:NPP_ < 1 for a clear majority of gridcells (IBIS, ISBA-CTRIP, JULES, LPX-Bern, VISIT-NIES). Several models exhibited multimodal distributions of *L*_Cveg*:NPP_ with one peak around zero, corresponding to no change in *C*_veg*_. No clear commonalities could be found among carbon-nitrogen coupled vs. C-only models (Fig. 2). On average, both groups exhibited a median of *L*_Cveg*:NPP_ very close to unity. Vegetation demography models overall have *L*_Cveg*:NPP_ >1, whereas non-demography models overall show *L*_Cveg*:NPP_ <1, inconsistent with theoretical expectation.

**Figure 3.**
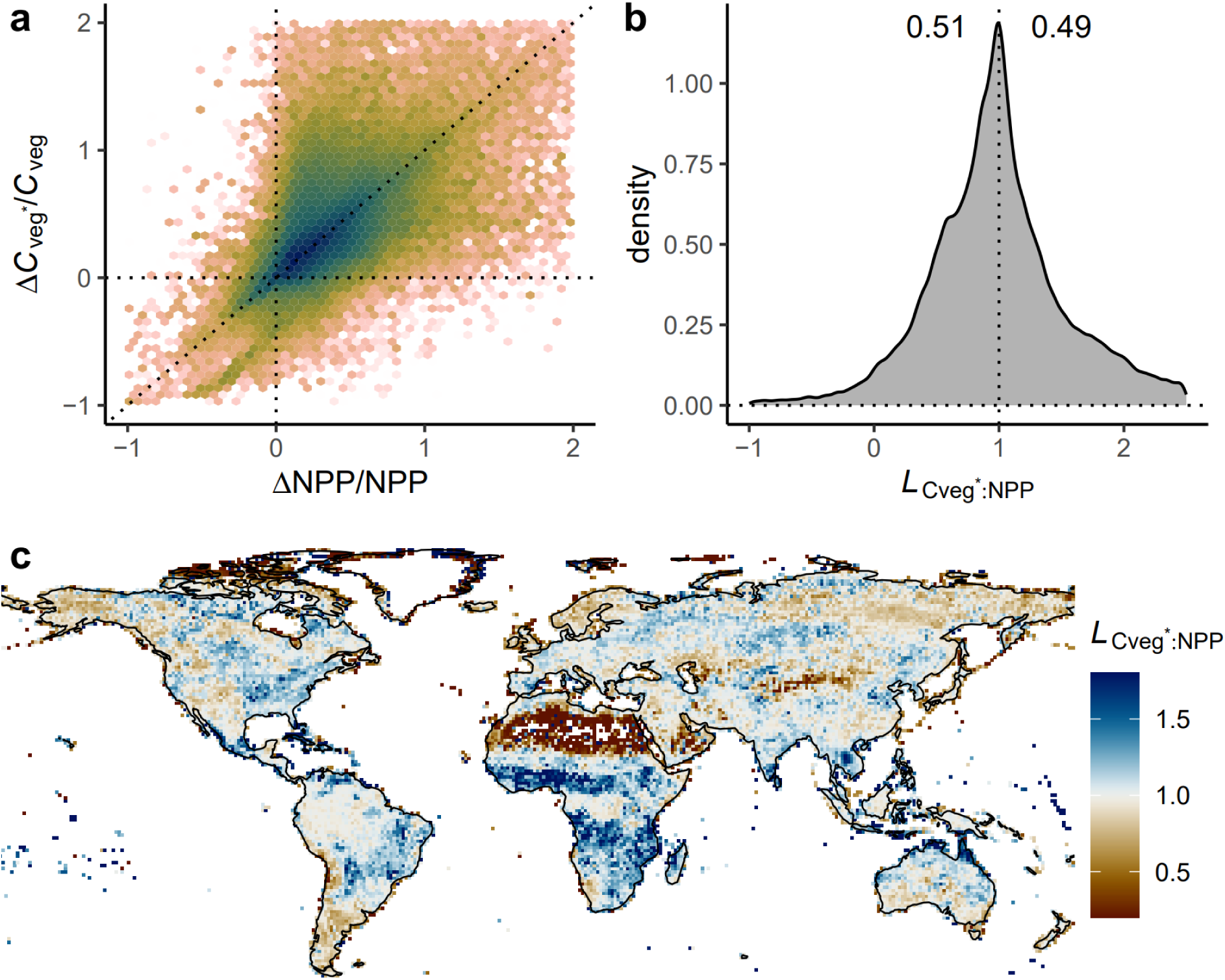
The relationship between the relative change of steady-state vegetation C (*R*Cveg* = ΔCveg*/Cveg) and the relative change of net primary productivity (*R*NPP = ΔNPP/NPP) under rising CO2 and changing N deposition, aggregated across models. (a) Density scatter plot of *R*Cveg* vs. *R*NPP values across all gridcells, pooled from balanced samples of all models. Dark colours denote high density, while bright colours denote low density. Values outside the axis ranges are not displayed. (b) Density of the distribution of *L*Cveg*:NPP (=*R*Cveg*/*R*NPP) values, pooled from balanced samples of all models. (c) Spatial pattern of *L*Cveg*:NPP, shown as the median per gridcell across models.

Geographical patterns emerge on average across all models, with a tendency for *L*_Cveg*:NPP_ < 1 in arid and cold regions (e.g., Sahara, northeastern Siberia, interior Eurasia) and *L*_Cveg*:NPP_ > 1 in seasonally dry tropical forests and savannah regions (Fig. 3c). However, spatial patterns of *L*_Cveg*:NPP_ are very diverse across individual models (Fig. S6). Across forested regions, several models consistently simulate *L*_Cveg*:NPP_ > 1 (CABLE-POP, CLASSIC, SDGVM), while other models consistently simulate *L*_Cveg*:NPP_ < 1 in forest regions (IBIS, ISAM, LPX-Bern). For several models (DLEM, ISAM, LPJ-GUESS, ORCHIDEE, VISIT, VISIT-NIES), *L*_Cveg*:NPP_ is near unity for tropical rainforests.

### The link between root and total vegetation carbon

Across models and gridcells, *L*_Croot:Cveg_ is distributed tightly around 1 (median 1.00, [IQR 0.32]), indicating overall a clear tendency of equal relative changes in root and total vegetation biomass and no general shift in allocation under rising CO_2_ (and changing N deposition) (Fig. 4). Likewise, *L*_Croot:Cveg_ tells a similar pattern to *L*_Croot:Cveg_. This pattern is also evident within most models and is most clearly expressed for CABLE-POP (Figs. S7 and S8). However, some models deviate from this pattern and simulate higher relative changes in root biomass than in total vegetation biomass (*L*_Croot:Cveg_ > 1) for a clear majority (>60%) of gridcells (IBIS, ISAM, VISIT-NIES). Some models simulate *L*_Croot:Cveg_ < 1 for a clear majority of gridcells (JULES, ORCHIDEE). The C-only models simulate, on average, a slightly higher *L*_Croot:Cveg_ (median 1.07) than C-N coupled models (median 1.0, Fig. 2). More allocation to *C*_root_ is often associated with less allocation to aboveground biomass (*C*_wood_ and/or *C*_leaf_). For *L*_Cleaf:Cveg*_, four models (CLM5.0 and LPX-BERN) show *L*_Cleaf:Cveg*_ > 1 while six models show very strong *L*_Cleaf:Cveg*_ < 1. Overall, the median of *L*_Cleaf:Cveg*_ is at 0.76 [IQR 0.81] suggesting less allocation to leaves under eCO_2_.

**Figure 4.**
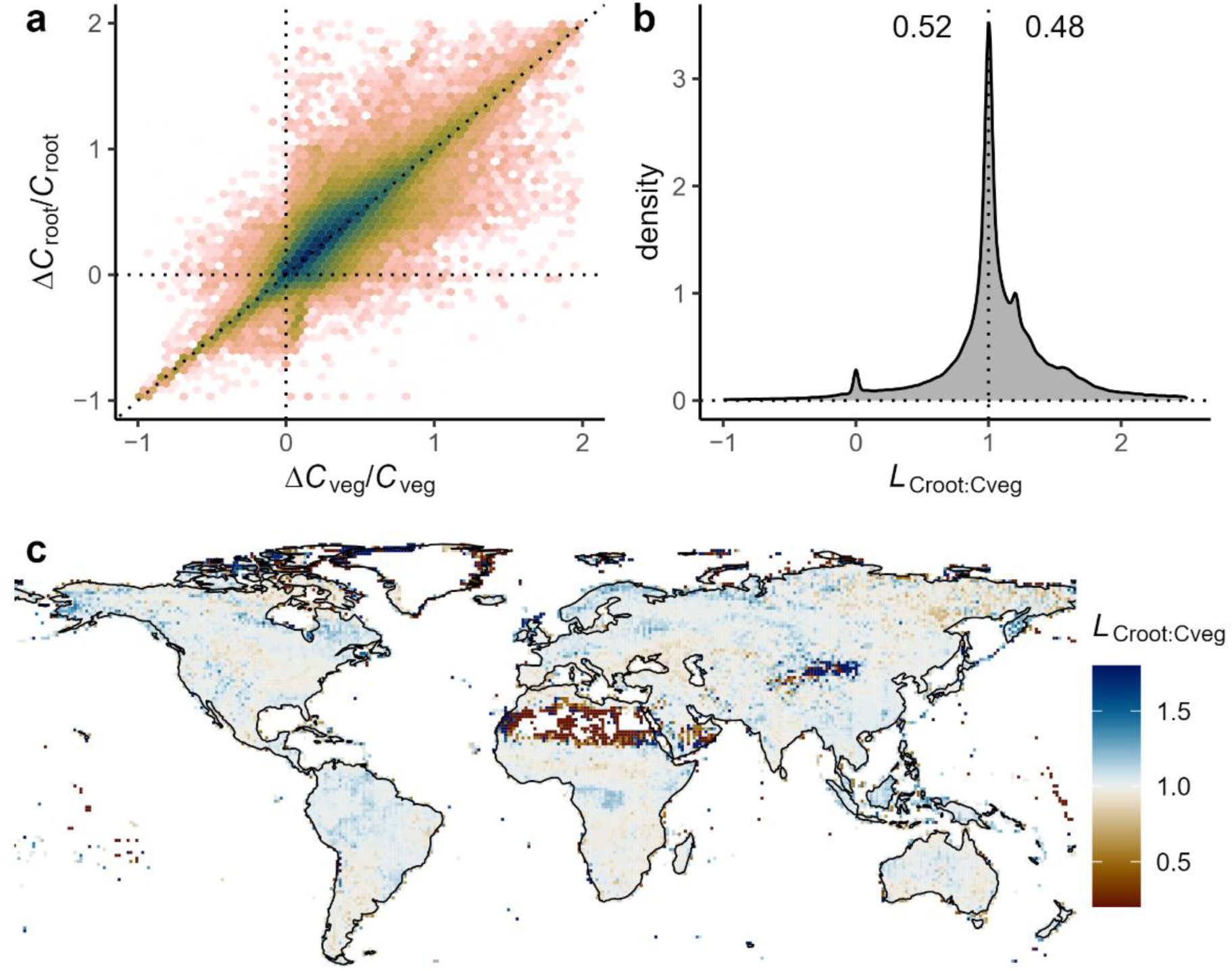
The relationship between the relative change of root biomass (*R*Croot = ΔCroot/Croot) and the relative change of total vegetation biomass (*R*Cveg = ΔCveg/Cveg) under rising CO2 and changing N deposition, aggregated across models. (a) Density scatter plot of *R*Croot vs. *R*Cveg values across all gridcells, pooled from balanced samples of all models. Dark colours denote high density, while bright colours denote low density. Values outside the axis ranges are not displayed. (b) Density of the distribution of *L*_Croot:Cveg_ (=*R*Croot / *R*Cveg) values, pooled from balanced samples of all models. (c) Spatial pattern of *L*_Croot:Cveg_, shown as the median per gridcell across models.

The geographical distribution of simulated *L*_Croot:Cveg_ varies across models (Fig. S9). For example, ORCHIDEE simulates values < 1 in high northern latitudes; JULES simulates values < 1 in boreal and tropical forest regions. IBIS and LPX-Bern simulate values > 1 in tropical forest regions, while ISAM and VISIT-NIES simulate values > 1 in most geographical regions. Some models exhibit a rather homogenous spatial pattern across the globe with values near 1 (CABLE-POP, CLM5.0, SDGVM), while others simulate strong spatial variations (LPX-Bern, CLASSIC).

## Discussion

We investigated the linearity of links along the modelled C cascade - from GPP to NPP, to vegetation biomass and its partitioning into different plant compartments. Our approach dissects multiple interacting processes and feedbacks along the C cascade into a set of diagnostically evaluable links that determine the response of the land C cycle to increasing CO_2_. Model structural choices have a strong influence on diagnosed *L* – as seen in the widely varying patterns in *L* for the different links and across the models analysed here. Comparison of diagnosed *L* against empirical patterns yields insight about model dynamics beyond commonly used static benchmarks and thereby tests model dynamics that are particularly relevant in the context of C cycle changes in response to the continued rise in atmospheric CO_2_.

*L* near unity has a clear interpretation: it indicates equal relative changes in two variables that are typically linked by a single or limited set of processes and model structural assumptions. *L* = 1 indicates that the simulated C cycle dynamics behave as a linear system and that “source-control” ^7,11,20,21^ dominates its response to increasing GPP under rising CO_2_. Thereby, diagnosing *L* distils key model behaviour that can be obtained from commonly available model outputs. Here, we used TRENDY simulations from CO_2_-only runs. *L* < 1 suggests an attenuating process along the C cascade – one that dampens the response of downstream pools or fluxes relative to upstream changes (e.g., in GPP). Conversely, *L* > 1 indicates an amplifying, or “super-proportional,” response, where downstream components of the C cycle respond more sensitively to changes in upstream C input.

We found widely varying patterns in *L* across models, in particular for the link between NPP and *C*_veg_. This indicates that alternative model structural choices lead to diverging model dynamics and that still no consensus exists about the representation of key mechanisms that govern the response of the land C cycle to rising CO_2_. This reflects findings from earlier works that evaluated a set of models against biomass measurements from FACE experiments (see a summary here ^57^). In contrast to these earlier works, our analysis covers more links and provides a global picture revealing that there is often a strong variation in model behaviour (measured by *L*) across the globe and different vegetation types and biomes – even within individual models. The clear deviation of *L* from unity for individual models and links also indicates that models embody process representations that imply an absence of purely source-driven dynamics.

Although models do not agree on the magnitude of the linearity (*L*) terms, deviations from unity were roughly balanced across the set of models investigated here. Hence, when analysed as the average across models, our analysis suggests a dominance of values near unity for each link and thus a tendency for an overall linear systems behaviour of terrestrial C cycle representations. Therefore, this set of models collectively simulates a linear propagation of CO_2_-driven GPP increases along the C cascade. That is, the relative enhancements of “downstream” fluxes and pools closely correspond to the relative enhancement of GPP under elevated CO_2_ (and changing N deposition). Previous empirical findings do point out that increases in GPP propagate across the C cascade and increase all “downstream” fluxes ^58^. However, links between fluxes reported by Litton et al. ^58^ were often not linear. Furthermore, departure from linear links between fluxes and pools could be driven by changes in effective turnover rates. Such non-linearities could arise, e.g., from the feedback of woody biomass production on respective turnover rates through the tree growth rate-longevity trade-off ^25–28^. Non-linearities may also result from changes in NPP partitioning, for example, a shift towards relatively more root production under increasing soil nutrient shortages ^17,59^, driven by their uptake and immobilisation through enhanced biomass production.

### GPP – NPP

The link between GPP and NPP is determined by the response of autotrophic respiration (*R*_a_) and the ecosystem carbon use efficiency (CUE = NPP/GPP) to rising CO_2_ and GPP. *R*_a_ is often conceived (and represented in models) as consisting of two components – growth and maintenance respiration ^60^. Growth respiration is commonly taken to be proportional to NPP, while the maintenance respiration is often taken to be controlled by the total mass of live biomass and modified by temperature and tissue nitrogen concentrations (LPX-Bern, LPJ-GUESS, CABLE-POP). In a few models (LPJ-GUESS, CABLE-POP), leaf maintenance respiration is modelled to be proportional to the capacity of Rubisco carboxylation in photosynthesis (*V*_cmax_) and (directly or indirectly) to leaf nitrogen content. This link to leaf N may introduce a feedback if leaf N concentrations are assumed to be affected by N availability. Hence, if N availability is simulated to decrease under rising CO_2_ – as is often the case in the absence of parallel increases in N deposition or experimental N fertilisation ^14,17^ – a reduction of leaf respiration may be simulated, leading to a CUE increase, and a pattern of *L*_NPP:GPP_ > 1. A similar feedback and resulting pattern of *L*_NPP:GPP_ > 1 may arise from CO_2_-driven reductions in *V*_cmax_ ^22^ if leaf respiration is formulated as a function of *V*_cmax_.

We found patterns of *L*_NPP:GPP_ > 1 across most gridcells for CLASSIC, JSBACH, LPJ. Among them, only JSBACH simulates C-N interactions, indicating that N limitation driving reduced leaf respiration may be an important process driving the results in JSBACH, but is not the dominating effect in all models exhibiting this pattern. It also indicates that reduced leaf respiration under elevated CO_2_ may also be simulated in C-only models (here, CLASSIC and LPJ). However, further investigations are needed to determine whether the observed pattern of *L*_NPP:GPP_ > 1 in several models is driven by changes in leaf respiration or other respiration components. Most C-only models show *L*_NPP:GPP_ > 1 except ISBA-CTRIP.

The simulated tendency for *L*_NPP:GPP_ < 1 in several models may be affected by the accounting of C components under N limitation. In C-N coupled models with widespread *L*_NPP:GPP_ < 1 (DLEM, LPX-Bern, ORCHIDEE, Fig. S1), other nitrogen limitation effects that increase autotrophic respiration may be the dominant driver of simulated responses to rising CO_2_. A mismatch of soil N supply and plant N demand is treated in some models by reducing NPP to match N supply under stoichiometric constraints and to divert C for N acquisition, whereby the invested C is accounted as an autotrophic respiration term, thus leading to reduced CUE (e.g. CLM5.0) ^61,62^. In other models, such C investments are counted towards allocation to a labile C pool and are thus part of NPP (CABLE-POP). Similarly, in JSBACH ^63^, NPP is defined before applying N limitation. After applying N limitation, part of NPP is allocated to labile pools, representing root exudates and other non-structural carbohydrates. In this study, JULES attributed such carbon to respiration not NPP, but Jones et al. ^64^ showed how JULES could be modified to simulate non-structural carbohydrates. This may contribute to the pattern of *L*_NPP:GPP_ > 1. For LPX-Bern, NPP is limited by the ratio of N demand to N uptake, while autotrophic respiration is not explicitly affected. Taken together, differences between models in their simulation of patterns in *R*_NPP_ vs. *R*_GPP_ are partly related to definition ambiguities in C components that are expended (as part of NPP) or respired (as part of *R*_a_) under N limitation. This results in a diversity of responses in *L*_NPP:GPP_ and its geographical pattern among C-N coupled models. Hence, a general behaviour of the simulated link between GPP and NPP could not be associated with whether models represented C-N interactions or not.

Observations from Free-Air CO_2_ Enrichment (FACE) experiments may provide the most direct test for simulated linearities diagnosed here. However, GPP is often not reported from FACE experiments because ecosystem-scale estimates are difficult to obtain. It is extremely challenging to distinguish measurements of NPP from biomass production (*C*_Veg_). Where GPP was estimated (EucFACE, POPFACE, DUKE), CUE increased and the relative increase in NPP tended to be higher than the relative increase in GPP ^24,65,66^. This appears consistent with the model ensemble that had *L*_NPP:GPP_ > 1. However, NPP increases in EucFACE, were likely predominantly driven by increased exudates and C export to mycorrhizae ^24^ – a component that is commonly not explicitly resolved in DGVMs. Plant biomass production increased much less than NPP in EucFACE. At DUKE FACE, measured biomass production increased by 25%, but Luo et al. ^65^ estimated a 40% increase in GPP by photosynthesis modelling. Increased C export to mycorrhizae under eCO_2_ is indeed widely reported ^67,68^. A revision of model output protocols and the provision of model results for biomass productivity (excluding allocation to non-structural C) would enable clearer comparability among models and against experimental results.

A widespread observation in experiments is that *V*_cmax_ declines under elevated CO_2_ ^17,69^ which implies a negative response of active Rubisco and may lead to a reduction in leaf respiration, contributing to an increase in CUE and *L*_NPP:GPP_ > 1 – consistent with simulations of several models analysed here and presented in the literature ^70–72^.

### NPP – C_veg*_

Two key mechanisms affect the link between NPP and *C*_veg_. First, *L*_Cveg*:NPP_ is affected by changes in the effective turnover time of C in biomass – a reflection of forest stand dynamics, tree demographic rates, self-thinning ^31^ and tree size-dependent mortality ^73^. Second, *L*_Cveg*:NPP_ is affected by changes in allocation of C to different pools with different turnover times – dominated by the distinction between allocation to woody vs. non-woody biomass ^35^. Thanks to the consideration of model outputs from CO_2_-only simulations, climate-driven changes in disturbance and environmental stress-related tree mortality are excluded by design. Yet, changes in tree mortality and effective C turnover times in biomass may still be affected indirectly by rising CO_2_ through accelerated self-thinning in vegetation demography-enabled models and other models that consider related processes through the crown area “packing constraint” ^31,74,75^. Moreover, shifts in allocation towards more short-lived fine root biomass under rising CO_2_ and altered nutrient availabilities could drive a reduction in effective turnover times in total vegetation C ^39^. Models did not report fine roots and coarse roots separately, which makes this difficult to diagnose. Nonetheless, we observed several models allocate considerably more to *C*_wood_ than *C*_leaf_ + *C*_root_ under eCO_2_ which implies an increase of the effective turnover times in *C*_veg_ (including JULES, CLASSIC, CABLE-POP and LPJ-GUESS) and thus theoretically a tendency for *L*_Cveg*:NPP_ >1.

Across models, the relationship between *R*_NPP_ and *R*_Cveg*_ was much weaker compared to the other two links investigated here. Yet, the ‘overall’ still indicates a tendency towards a predominantly linear relationship with the distribution of *L*_Cveg*:NPP_ peaking around 1.0 (Fig. 2d). This indicates that, across this model ensemble, a change in NPP elicits a relatively wide range of changes in steady-state *C*_veg_, but on average across the globe and the model ensemble, a linear relationship dominates. A secondary peak of the *L*_Cveg*:NPP_ distribution around 0 (Fig. S20) indicates that no vegetation biomass changes are simulated despite NPP increases. This pattern occurs mostly in arid and grassland regions (Fig. S6) (Fig. 3 compared to Fig. S20) and likely reflects the behaviour of annual herbaceous vegetation in which NPP increases do not lead to sustained *C*_veg_ gains.

For several models, we found *L*_Cveg*:NPP_ around unity or slightly above across forest regions (ISBA-CTRIP, ORCHIDEE, VISIT, VISIT-NIES), indicating no or compensating responses in effective biomass turnover rates and mortality and allocation to short vs. long-lived biomass pools (Fig. S6). Several models that consider a “packing constraint” for simulating tree size-density relationships (LPJ, LPX-Bern) exhibit general patterns of *L*_Cveg*:NPP_ < 1 across tropical forest regions (Fig. S6). In LPJ-GUESS, this packing constraint is partly relieved by allowing tree crown overlaps ^75^, which may result in higher *L*_Cveg*:NPP_ values than in the genealogically related models LPJ and LPX-Bern. In IBIS, a general pattern of *L*_Cveg*:NPP_ < 1 may be linked to a smaller increase in woody biomass than in total biomass (Fig. 2). Simulated patterns of *L*_Cveg*:NPP_ < 1 may also arise from structural limits of tree sizes and ensuing mortality that are reached more often as NPP increases (JSBACH). In contrast, SDGVM assumes a positive relationship between the wood allocation fraction and NPP ^75^ and exhibits a general pattern with *L*_Cveg*:NPP_ > 1 across forest regions. Additionally, CABLE-POP simulates more carbon allocation to total vegetation than to leaves (Fig. 2) and exhibits a dominant *L*_Cveg*:NPP_ > 1. As explained here, general patterns of *L*_Cveg*:NPP_ are linked to model structural assumptions. Nonetheless, other mechanisms could be at play and simulated patterns are likely affected by simulated vegetation states (e.g., tree density and size). Therefore, direct causal effects between model structure and diagnosed linearity could be better interpreted if additional model outputs were made available (e.g. NPP of vegetation component; Tree mortality).

It is challenging to assess how well the simulated patterns align with empirical evidence for *L*_Cveg*:NPP_. A pattern of *L*_Cveg*:NPP_ < 1 in forest regions appears to be supported by the observation that fast-growing trees tend to live shorter than slow-growing trees ^25,27–29^. However, these relationships predominantly emerge from spatial or interspecific variations ^8,29^ not from temporal changes in response to eCO_2_. Yet, one long-term forest monitoring study found little change in forest self-thinning relations despite accelerated growth, thus supporting *L*_Cveg*:NPP_ < 1 ^76^. In contrast, an analysis of FACE experiments in relatively young forests indicates increases in wood allocation with increasing NPP ^35^, thus supporting *L*_Cveg*:NPP_ > 1. Nonetheless vegetation demography models overall have *L*_Cveg*:NPP_ >1, against the theoretical expectation. The application of vegetation demography should reduce *L*_Cveg*:NPP_, but this signal may be masked by other processes, for example, an increase in *C*_veg_ turnover time due to more allocation to woods than leaf (Fig. 2).

Future studies are needed to investigate the relative contribution of different mechanisms that affect the link between NPP and the vegetation C pool and to use diverse observations for estimating linearity in the framework applied here.

### C_veg_ – C_root_

The link between *C*_veg_ and *C*_root_ is determined by allocation and whether the effective turnover rates in these pools change under rising CO_2_. Unless effective turnover rates change, an increased fraction of NPP allocated to root biomass will yield *L*_Croot:Cveg_ > 1. There is evidence from ecosystem experiments for such a shift towards more belowground allocation under elevated CO_2_ ^22,24,36–41^ and that the relative increase in *C*_root_ is greater than the relative increase in *C*_veg_ ^17^. This response has been interpreted in light of the functional balance hypothesis ^77–79^ which predicts a shift towards more belowground allocation as the balance between CO_2_ assimilation efficiency (C uptake per unit aboveground biomass) and soil nutrient acquisition efficiency (nutrient acquisition per unit root biomass) increases under rising CO_2_ ^17^. However, dynamics in response to an experimental step-change in CO_2_ likely differ from dynamics in response to a gradual rise, extended over decadal to centennial time scales as simulated in TRENDY runs. In models that simulate a demand-dependency of N uptake, previous multi-model evaluations against FACE experiments found a gradual relaxation from N limitation over time ^75^. Hence, the gradual rise in CO_2_ may trigger a weaker nitrogen limitation effect and associated shifts in allocation, and a reduced tendency for *L*_Croot:Cveg_ > 1.

On average across the ensemble of models investigated here, this pattern towards increased belowground allocation is not generally simulated. Contrary to expectations, C–N models simulate lower *L*_Croot:Cveg_ than C-only models. Only ISAM, IBIS, and VISIT-NIES simulated a clear tendency for *L*_Croot:Cveg_ > 1. ISAM accounts for C-N cycling and a plastic response of root growth to soil N supply and demand ^80^, which likely led to the clear pattern of *L*_Croot:Cveg_ > 1. However, other models that exhibited *L*_Croot:Cveg_ > 1 (IBIS, VISIT-NIES) did not resolve N cycling, and models that did resolve C-N interactions simulated *L*_Croot:Cveg_ < 1 for a clear majority of gridcells (JULES, ORCHIDEE). Several models simulated predominantly equal relative changes in total vegetation biomass and root biomass. The distribution of *L*_Croot:Cveg_ is most narrowly constrained around 1 for IBIS, CABLE-POP and SDGVM, and ORCHIDEE (shown as a tall peak in Fig.S7 and Fig.S16). This pattern likely reflects the most common model structural choice whereby fixed allocation is considered per plant functional type. E.g., ORCHIDEE considers a fixed stem to coarse root ratio following the ‘pipe model’ concept ^81^. Note that *C*_root_ of ORCHIDEE includes coarse roots (Fig. S19). CABLE-POP considers a fixed allocation fraction to fine roots ^82^ (Fig. S7). In IBIS, the distribution of *L*_Croot:Cveg_ exhibits two modes and it appears that root allocation in forest regions is governed differently, leading to *L*_Croot:Cveg_ > 1, compared to other regions, where *L*_Croot:Cveg_ is near unity (Fig. S9). Substantial between-model structural differences exist also regarding allocation to *C*_leaf_ and *C*_wood_ (Figs. S11 to S16). We note that the interpretation of *L*_Croot:Cveg_ patterns in relation to changes in root allocation should be approached with caution, as concurrent shifts in effective turnover rates can obscure or counteract these patterns. Additional model output variables, most importantly plant compartment-specific NPP, would enable a clearer interpretation of simulated mechanisms.

### Limitations

Diagnosing linearity in responses of the land C cycle is challenged by the fact that available simulation outputs do not represent steady states under different levels of CO_2_ but instead reflect the transient response. We applied an estimation of the steady state from transient simulations (Eq. 12). In future, purposely designed model simulations (e.g., a step-increase in CO_2_ and a subsequent equilibration to a new steady-state), or model outputs on plant compartment NPP would facilitate diagnosing system dynamics and the comparison to ecosystem CO_2_ experiments. Our approach to diagnosing system dynamics from available transient model outputs enables comparisons to diverse observations in the future and may guide future model development and evaluation towards a focus on capturing essential and widely observed patterns in the C cycle dynamics. Currently, we explained theoretical expectations of *L*_NPP:GPP_ >1 and *L*_Cveg*:NPP_ <1. However, a data-model comparison for these two links is not possible owing to the lack of field data. Considering the difficulties in measuring GPP and NPP (beyond C_veg_), data availability is not likely to drastically change in near future. Both empirical and modelling studies are suggested to benchmark individual process behind the two links, for example the effect of eCO_2_ on root exudates, self-thinning and more as discussed above. For *L*_Croot:Cveg_, empirical evidence is ample, albeit not consistent. Nonetheless, if coarse root and fine root NPP are provided by models in the future, data-model comparison for this link may become possible.

Furthermore, for the *L* terms quantified on links that include pools (here *L*_Cveg*:NPP_ and *L*_Croot:Cveg*_), interpretations have to consider that both changes in relative allocation between compartments with widely varying turnover rates and compartment-specific turnover rates (affected by continuous turnover and by more episodic tree mortality, phenology and leaf senescence) can influence results. This obscures interpretations with a direct view to process representations in the models. Additional model outputs (e.g., compartment-specific NPP) could be used to better inform analyses in this respect.

### Conclusions

Overall, we found that for the model ensemble, a linear behaviour dominates, whereby the relative change in GPP largely dictates relative changes in downstream fluxes (NPP) and pools. This linearity implies that, collectively, DGVMs still largely adhere to a source-driven conceptualisation of terrestrial C dynamics. However, individual models clearly deviate from linear dynamics. Our analysis did not yield any support that this deviation from linear dynamics is a reflection of (more recently added) representations of sink controls (e.g., N limitation), and vegetation demography. We found the strongest inter-model disagreement for the link between vegetation C and NPP – indicating a need for model benchmarking with a focus on related processes – including tree growth-biomass relationships and forest demographic processes, CO_2_-driven changes in allocation to tissues with widely differing turnover times.

## Acknowledgements

H.Z.-Z. was supported as part of the Next Generation Ecosystem Experiments-Tropics, funded by the U.S. Department of Energy, Office of Science, Office of Biological and Environmental Research. JSBACH simulations are performed using resources of the Deutsches Klimarechenzentrum (DKRZ) granted by its Scientific Steering Committee (WLA) under project 891. A.I. was supported by JSPS KAKENHI (grant no. 21H05318). Q.S. was supported by the Swiss National Science Foundation (200020 200511). Simulations of LPX-Bern were performed on UBELIX (https://www.id.unibe.ch/hpc), the HPC cluster at the University of Bern. T.A.M.P. was funded under the European Union’s Horizon 2020 programme (grant agreement no. 758873, TreeMort) and Horizon Europe (101141836, Tree2Globe). This study is a contribution to the Swedish government’s strategic research areas BECC and MERGE and the Nature-based Future Solutions profile area at Lund University. ORNL is managed by UT-Battelle, LLC, for the DOE under contract DE-AC05-1008 00OR22725. BDS acknowledges funding from the Swiss National Science Foundation grant PCEFP2_181115.

## Authors Contribution

B.D.S conceived the study. H.Z.-Z., B.D.S., R.D., and M.T. carried out data analysis. H.Z.-Z. and B.D.S wrote the manuscript with inputs and revision from Q.S., V.A., P.A., T.A.M.P., A.K.J., W.Y., J.N., D.S.G., J.P., B.P., A.P.W., S.Z., P.B., J.K., E.K., S.S., M.O., H.T., N.P., P.F., A.I., and J.D. ran the models and provided model outputs.

## Code and data availability

Codes and data to reproduce figures are currently available on <u>GitHub</u>. The same repository is uploaded to ‘figshare’ during submission. To reproduce results, please run vignettes /DGVMLIN_trendy_v11.rmd. Values for the *R* and *L* terms are available as **<u>supplementary data</u>**.

## 1. Linearity *L*_NPP:GPP_

**Figure S1.**
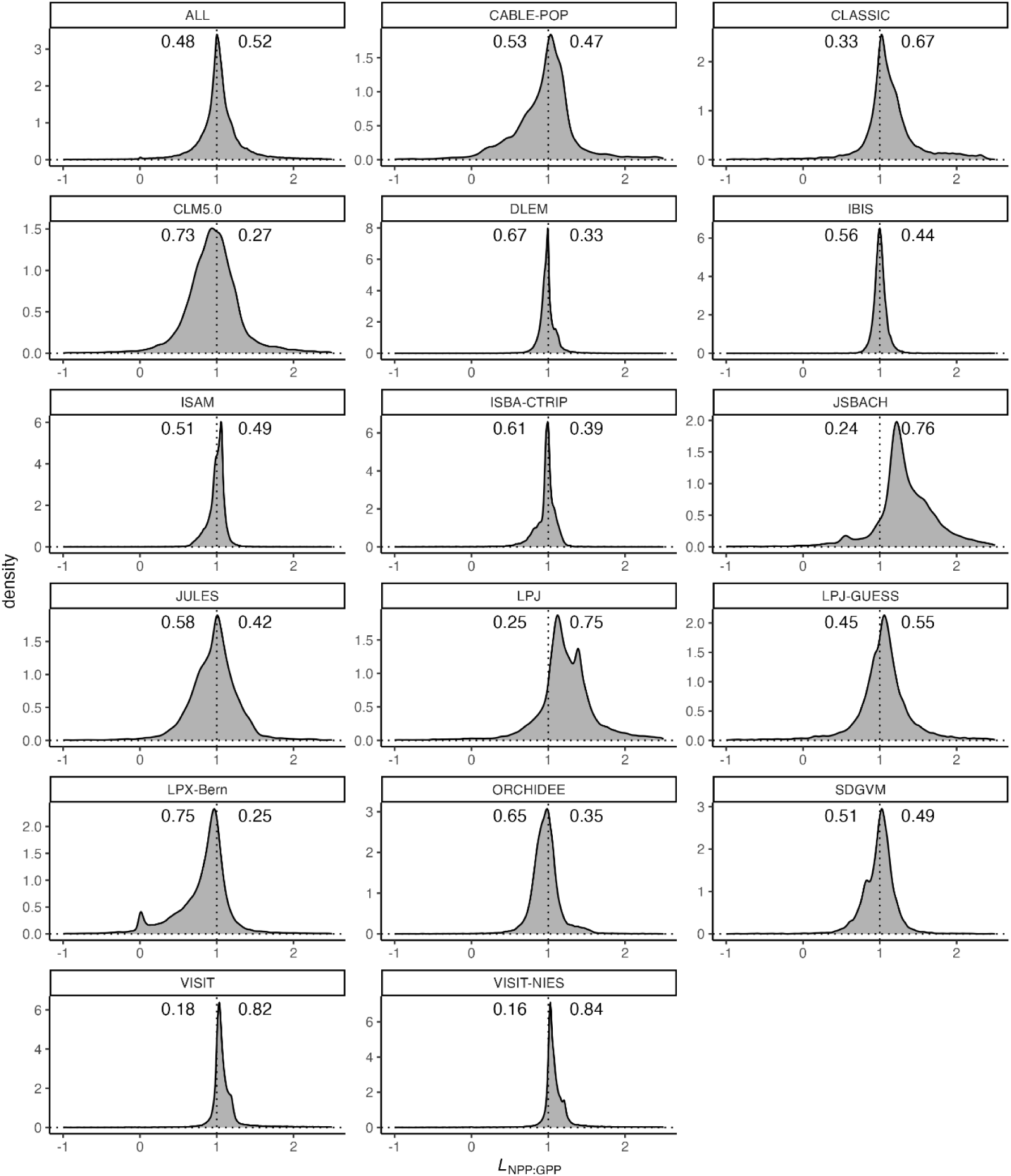
Density of the distribution of *L*_NPP:GPP_ values across gridcells by model, printed as labels on top of each panel. ‘All’ shows cross models overall pattern, identical to the figure presented in the main text. The proportion of gridcells with *L*_NPP:GPP_ < 1 (*L*_NPP:GPP_ > 1) is given by the annotation on the left (right) side of the plotting area. The panel labelled ‘ALL’ represents the joint pattern with data pooled from all models. *L*_NPP:GPP_ = *R*_NPP_ / *R*_GPP_ where *R*_NPP_ (*R*_GPP_) is the relative change of net primary productivity, NPP (gross primary production, GPP), evaluated from simulations with rising CO_2_ and changing N deposition. Both *L* and *R* terms are unitless.

**Figure S2.**
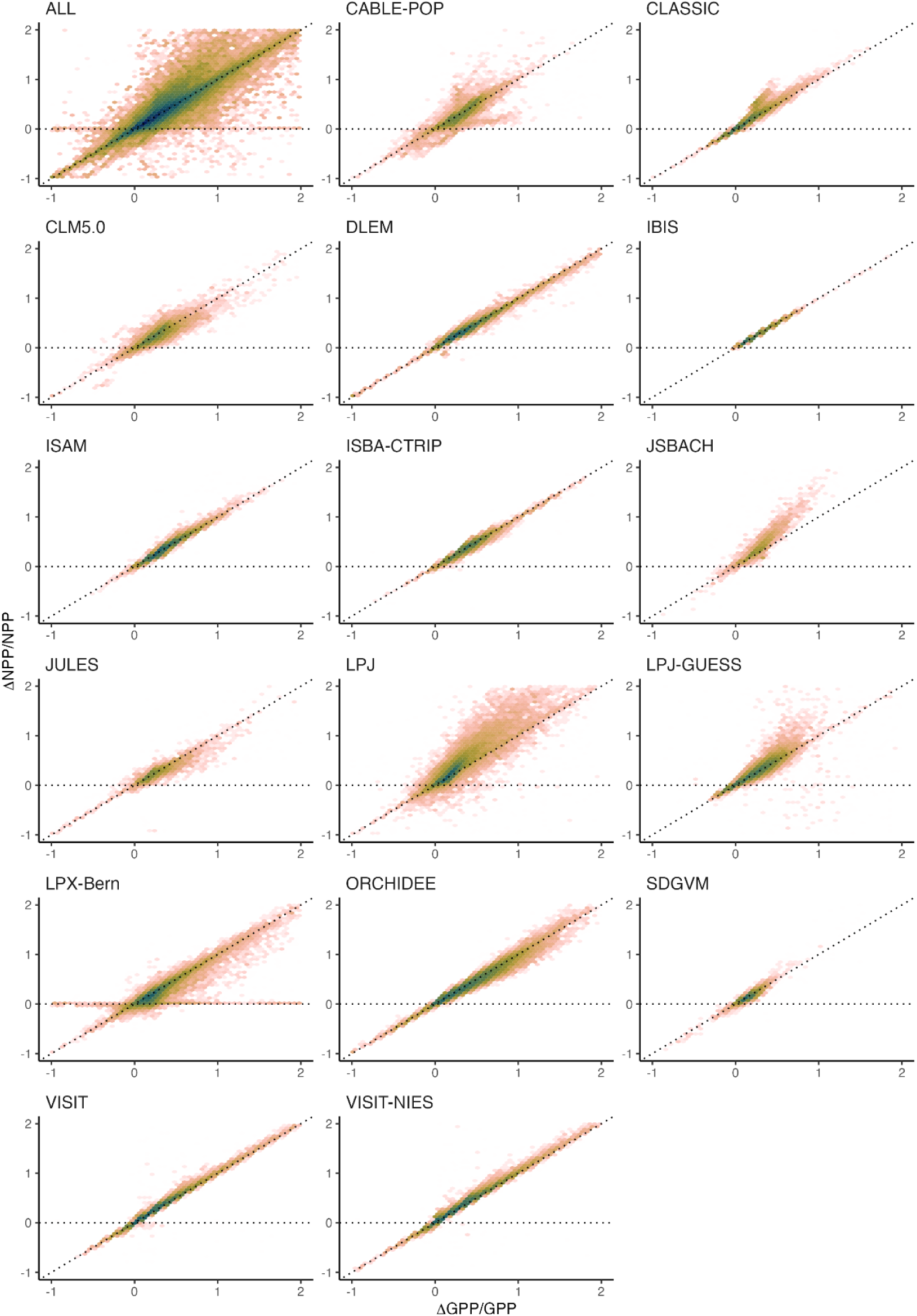
Density scatter plot of *R*_NPP_ vs. *R_G_*_PP_ values across all gridcells for each model. The name of the model is printed on top of each histogram. Dark colors denote high density, while bright colors denote low density. *R*_NPP_ is the relative change (ΔNPP/NPP, unitless) of net primary productivity (NPP) evaluated from simulations with rising CO_2_ and changing N deposition. *R*_GPP_ is the relative change (ΔGPP/GPP, unitless) of gross primary productivity (GPP), evaluated from the same simulations. The panel labelled ‘ALL’ represents the joint pattern with data pooled from all models.

**Figure S3.**
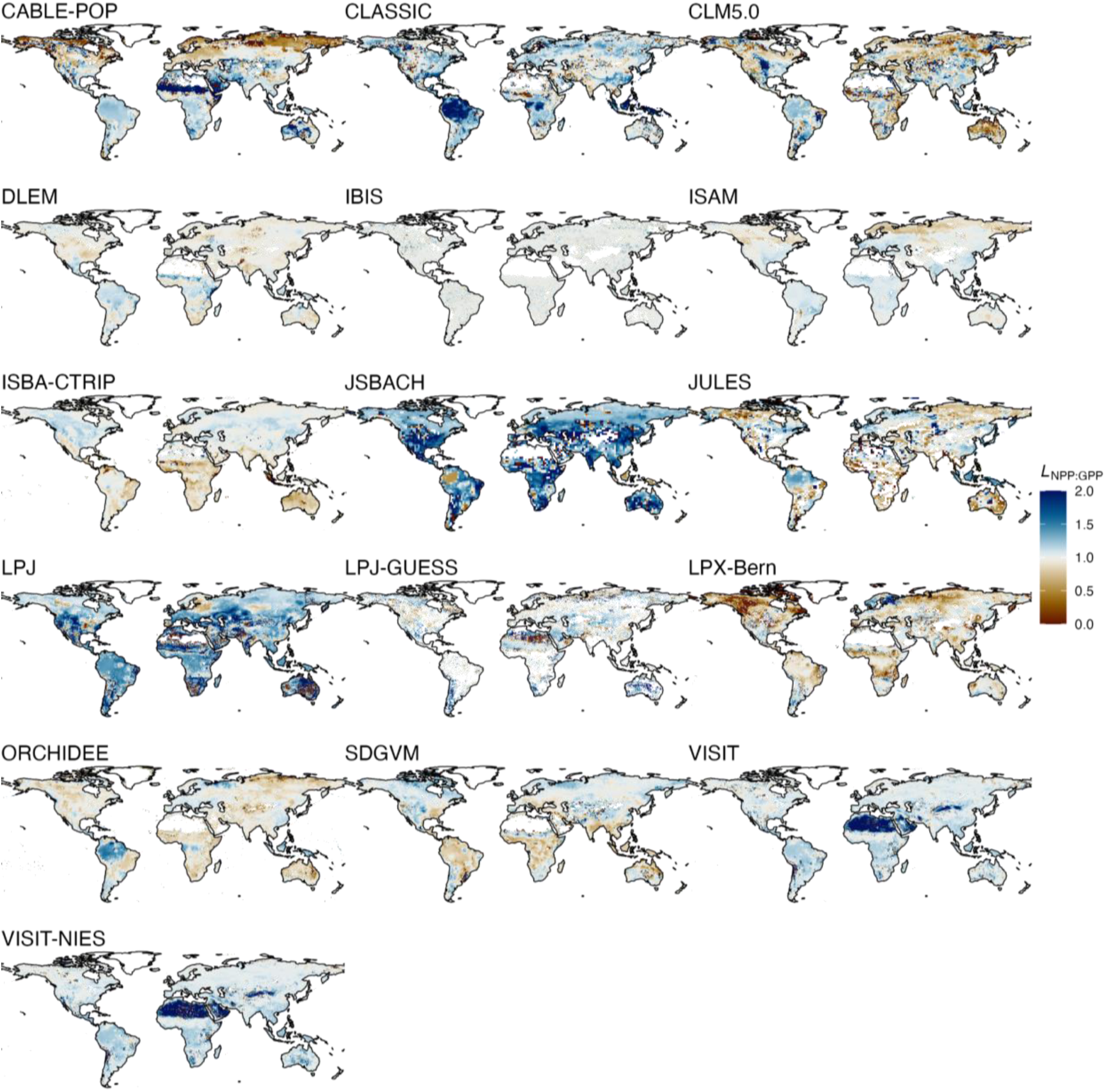
Spatial pattern of *L*_NPP:GPP_ for each model. The name of each model is printed on top of each map. *L*_NPP:GPP_ = *R*_NPP_ / *R*_GPP_ where *R*_NPP_ (*R*_GPP_) is the relative change of net primary productivity, NPP (gross primary production, GPP), evaluated from simulations with rising CO_2_ and changing N deposition. Both *L* and *R* terms are unitless.

## 2. Linearity *L*_Cveg*:NPP_

**Figure S4.**
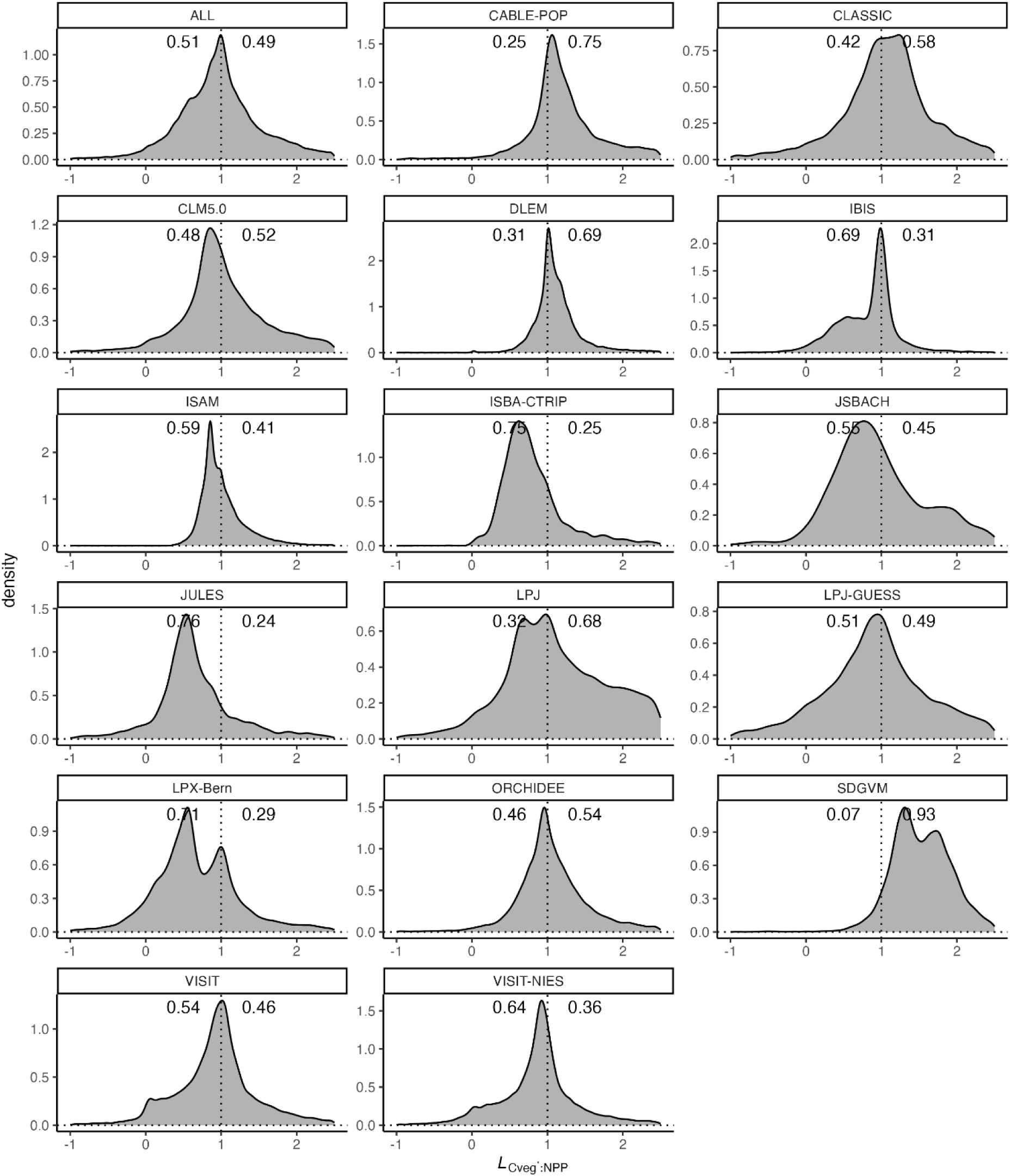
Density of the distribution of *L_Cveg*:NPP_* values across gridcells by model, printed as labels on top of each panel. The proportion of gridcells with *L*_NPP:GPP_ < 1 (*L*_NPP:GPP_ > 1) is given by the annotation on the left (right) side of the plotting area. The panel labelled ‘ALL’ represents the joint pattern with data pooled from all models. *L*_Cveg*:NPP_ = *R*_Cveg*_ / *R*_NPP_ where *R*_NPP_ (*R*_Cveg_) is the relative change of net primary productivity, NPP (steady-state total vegetation C pools), evaluated from simulations with rising CO_2_ and changing N deposition. Both *L* and *R* terms are unitless.

**Figure S5.**
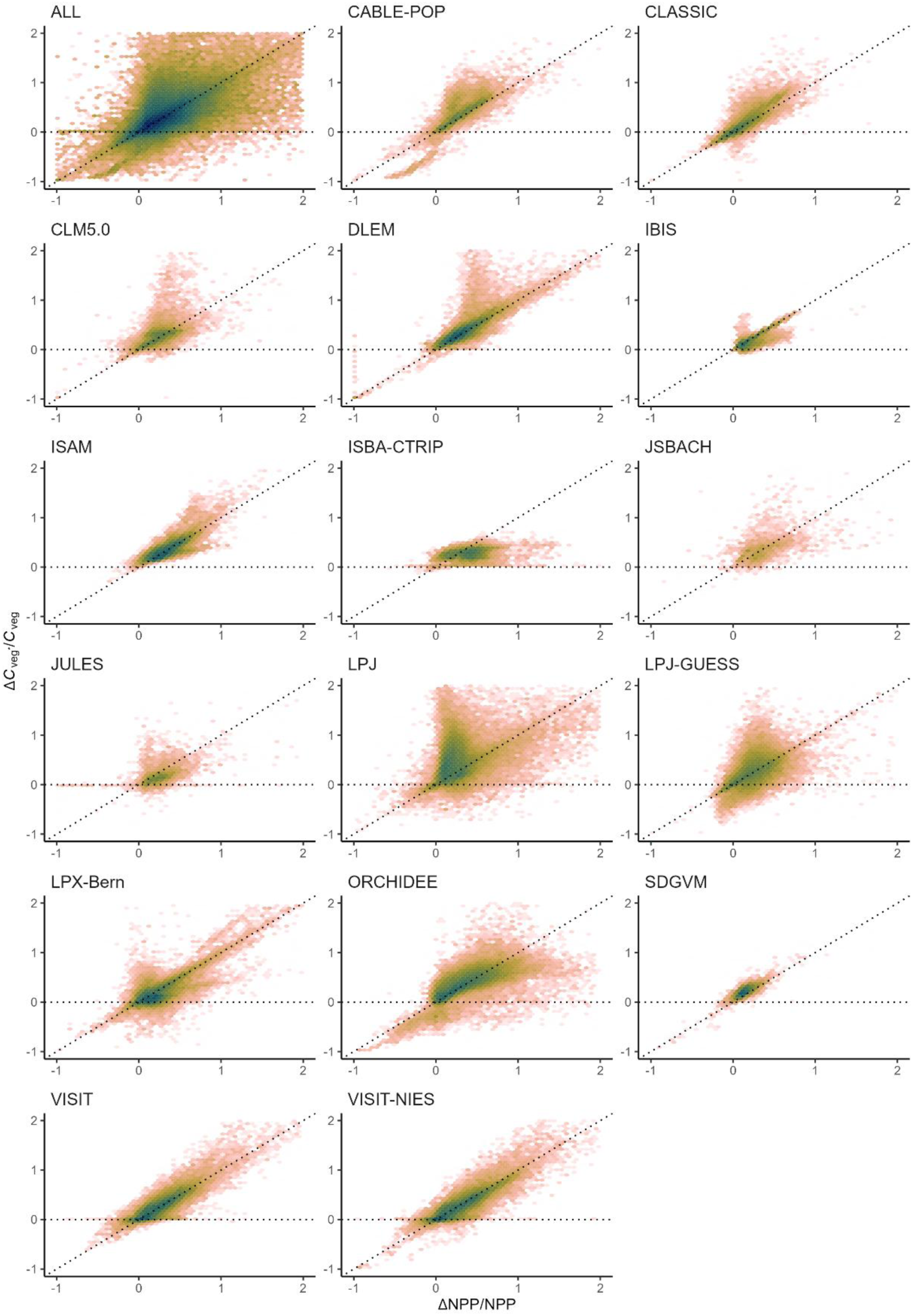
Density scatter plot of *R*_NPP_ vs. *R*_Cveg*_ values across all gridcells for each model. The name of the model is printed on top of each histogram. Dark colors denote high density, while bright colors denote low density. *R*_NPP_ is the relative change (ΔNPP/NPP, unitless) of net primary productivity (NPP) evaluated from simulations with rising CO_2_ and changing N deposition. *R*_Cveg*_ is the relative change (ΔC_veg*_/C_veg*_, unitless) of total vegetation carbon, evaluated from the same simulations. The panel labelled ‘ALL’ represents the joint pattern with data pooled from all models.

**Figure S6.**
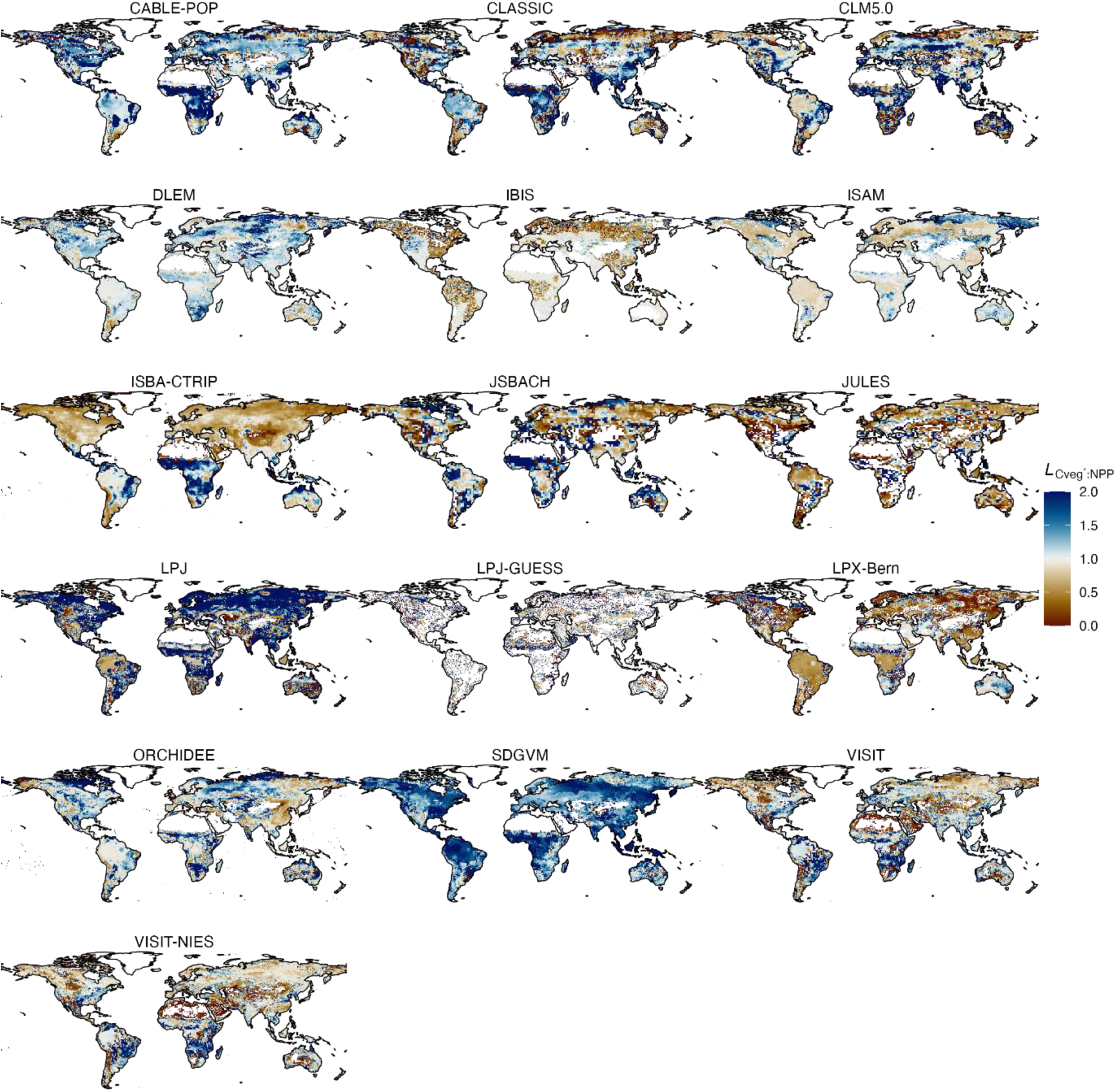
Spatial pattern of *L*_Cveg*:NPP_ for each model. The name of each model is printed on top of each map. *L*_Cveg*:NPP_ = *R*_Cveg*_ / *R*_NPP_ where *R*_Cveg*_ is the relative change of the steady-state estimate of total vegetation carbon, and *R*_NPP_ is the relative change in net primary production (NPP), evaluated from simulations with rising CO_2_ and changing N deposition. Both *L* and *R* terms are unitless.

## 3. Linearity *L*_Croot:Cveg_

**Figure S7.**
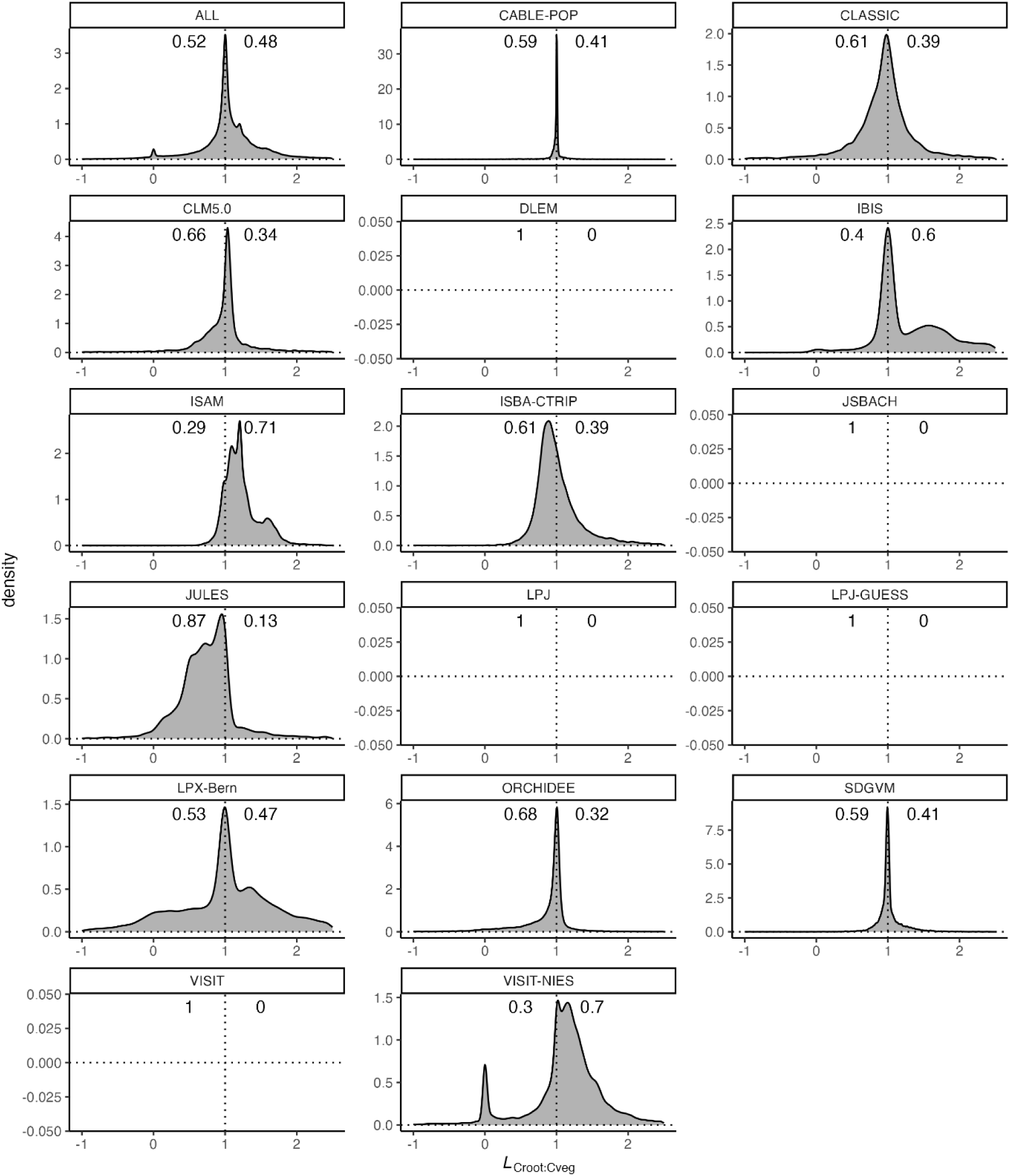
Density of the distribution of *L*_Croot:Cveg_ values across gridcells by model, printed as labels on top of each panel. The proportion of gridcells with *L*_Croot:Cveg_ < 1 (*L*_Croot:Cveg_ > 1) is given by the annotation on the left (right) side of the plotting area. The panel labelled ‘ALL’ represents the joint pattern with data pooled from all models. *L*_Croot:Cveg_ = *R*_Croot_ / *R*_Cveg_ where *R*_Croot_ (*R*_Cveg_) are the relative change of root C (total vegetation C), evaluated from simulations with rising CO_2_ and changing N deposition. Both *L* and *R* terms are unitless.

**Figure S8.**
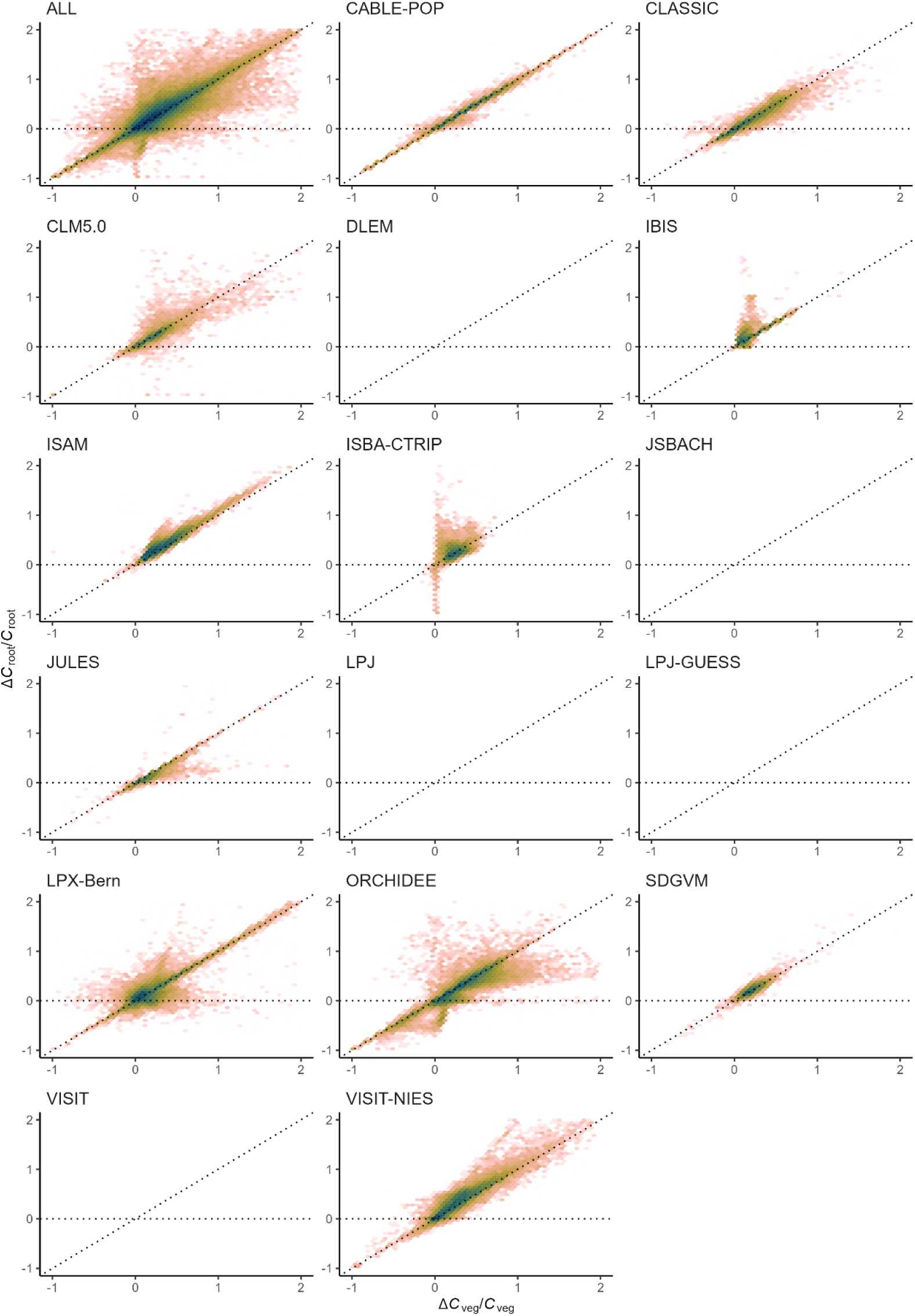
Density scatter plot of *R*_Croot_ vs. *R*_Cveg_ values across all gridcells for each model. The name of the model is printed on top of each histogram. Dark colors denote high density, while bright colors denote low density. *R*_Croot_ is the relative change (ΔC_root_/C_root_, unitless) of root C. *R*_Cveg_ is the relative change (ΔC_veg_/C_veg_, unitless) of total vegetation C. Both relative changes are evaluated from the same simulations. The panel labelled ‘ALL’ represents the joint pattern with data pooled from all models.

**Figure S9.**
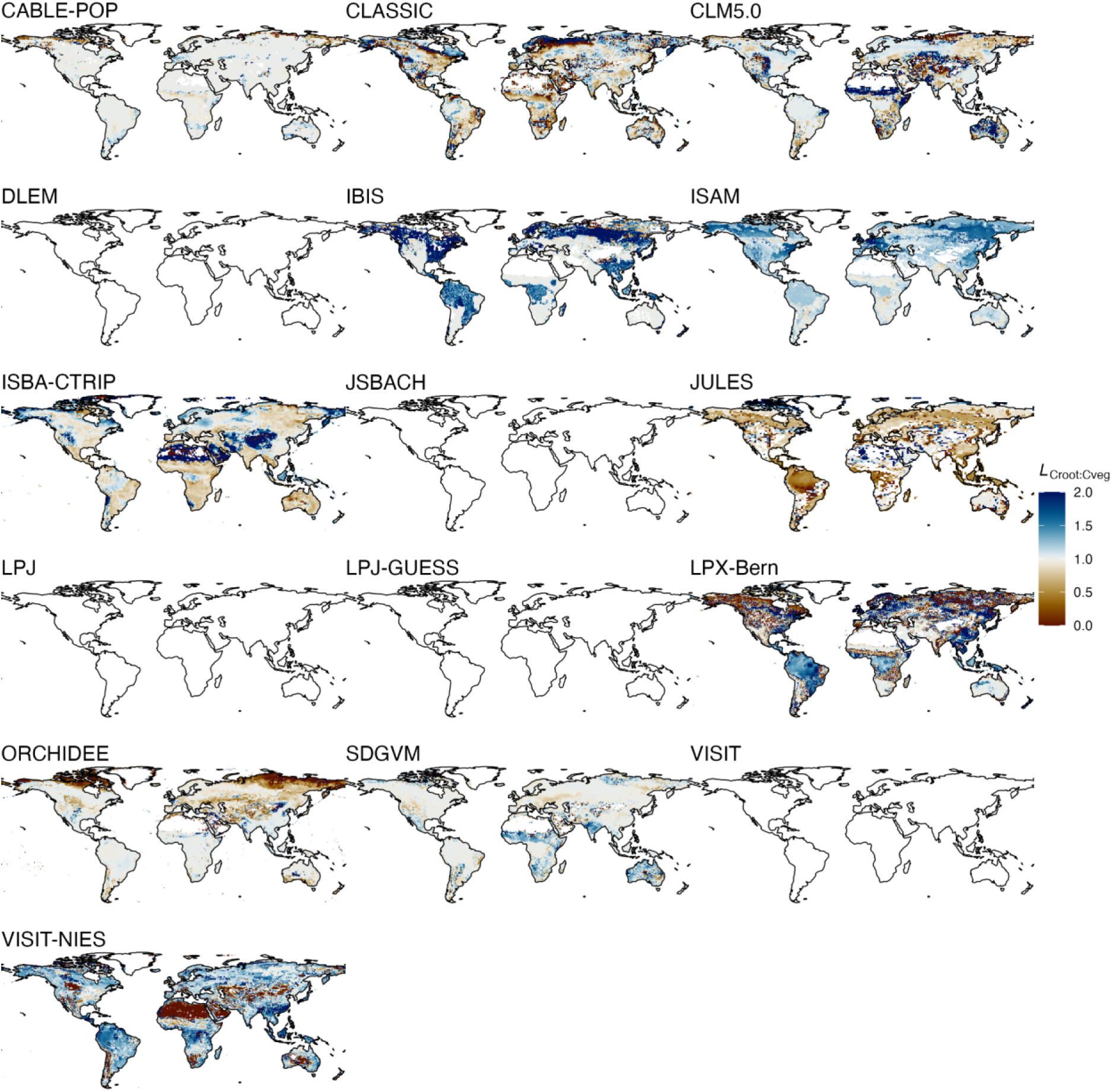
Spatial pattern of *L*_Croot:Cveg_ for each model. The name of each model is printed on top of each map. *L*_Croot:Cveg_ = *R*_Croot_ / *R*_Cveg_ where *R*_Croot_ (*R*_Cveg_) is the relative change in root C (total vegetation C), evaluated from simulations with rising CO_2_ and changing N deposition. Both *L* and *R* terms are unitless.

## 4. Linearity *L*_Croot:Cveg*_

**Figure S10.**
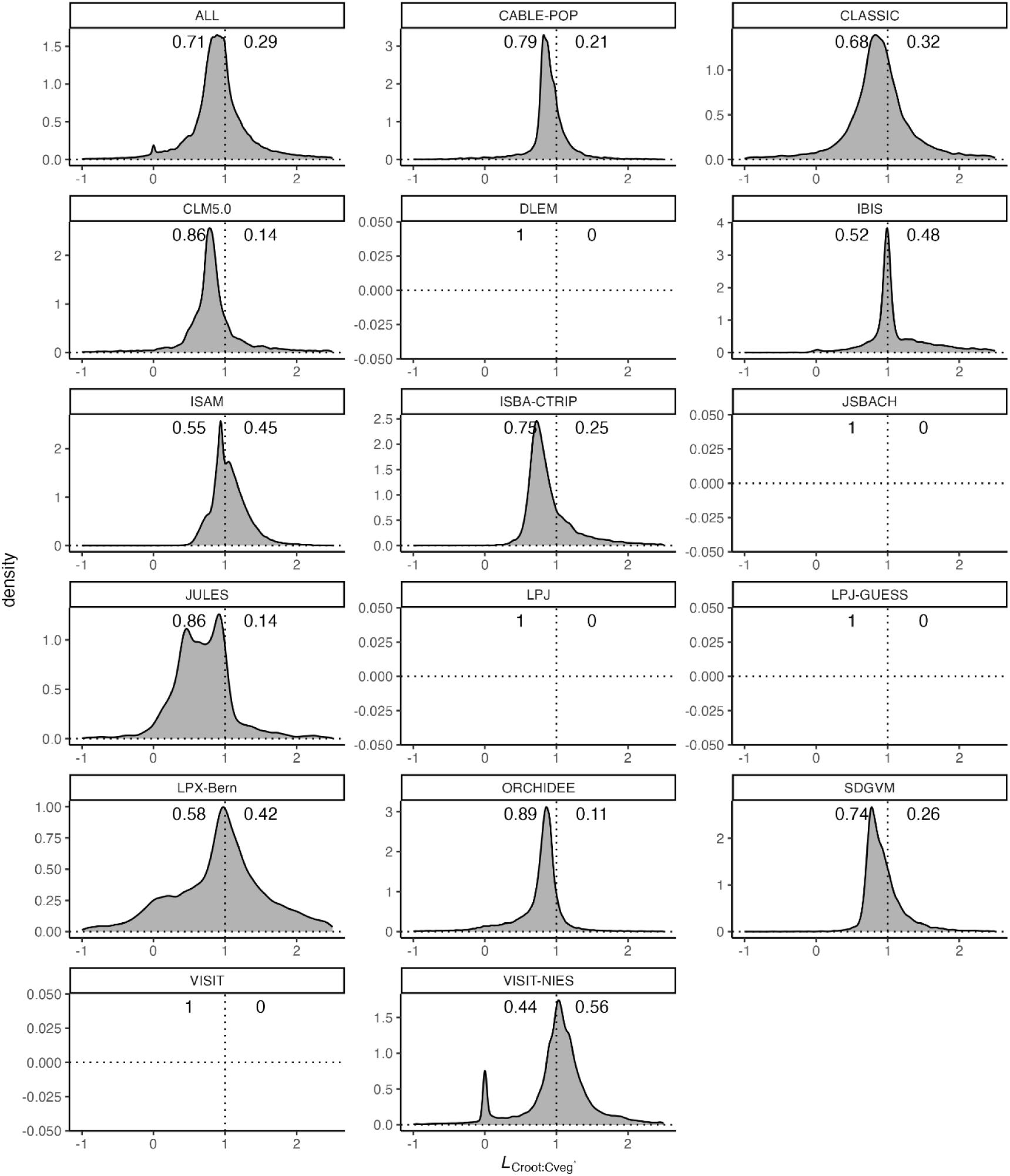
Density of the distribution of *L*_Croot:Cveg*_ values across gridcells by model, printed as labels on top of each panel. The proportion of gridcells with *L*_Croot:Cveg*_ < 1 (*L*_Croot:Cveg*_ > 1) is given by the annotation on the left (right) side of the plotting area. The panel labelled ‘ALL’ represents the joint pattern with data pooled from all models. *L*_Croot:Cveg*_ = *R*_Croot_ / *R*_Cveg*_ where *R*_Croot_ are the relative change of root C (steady-state total vegetation C), evaluated from simulations with rising CO_2_ and changing N deposition. Both *L* and *R* terms are unitless. As many models include coarse roots in *C*_root_, *C*_root_ should not have rapid response to carbon fertilization. Ideally, we should apply steady state conversion to *C*_root_, and calculate *L*_Croot*:Cveg*_ but this could not be implemented due to the lack of *C*_root_ turnover time. We believe that *L*_Croot:Cveg*_ is less useful than *L*_Croot:Cveg_.

**Figure S11.**
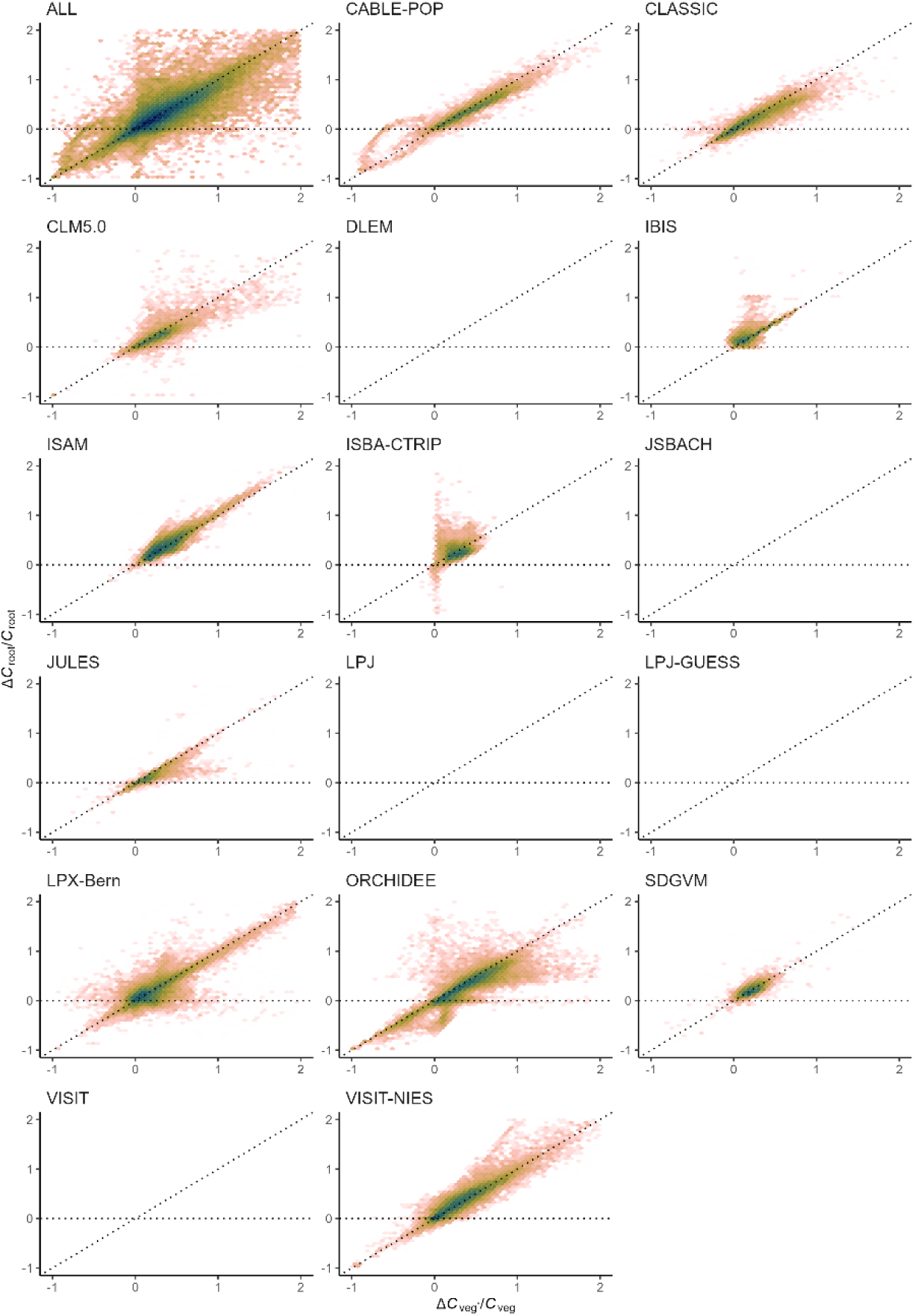
Density scatter plot of *R*_Croot_ vs. *R*_Cveg*_ values across all gridcells for each model. The name of the model is printed on top of each histogram. Dark colors denote high density, while bright colors denote low density. *R*_Croot_ is the relative change (ΔC_root_/C_root_, unitless) of root C. *R*_Cveg*_ is the relative change (ΔC_veg*_/C_veg_, unitless) of steady-state total vegetation C. Both relative changes are evaluated from the same simulations. The panel labelled ‘ALL’ represents the joint pattern with data pooled from all models. As many models include coarse roots in *C*_root_, *C*_root_ should not have rapid response to carbon fertilization. Ideally, we should apply steady state conversion to *C*_root_, and calculate *L*_Croot*:Cveg*_ but this could not be implemented due to the lack of *C*_root_ turnover time. We believe that *L*_Croot:Cveg*_ is less useful than *L*_Croot:Cveg_ (Figure S8).

**Figure S12.**
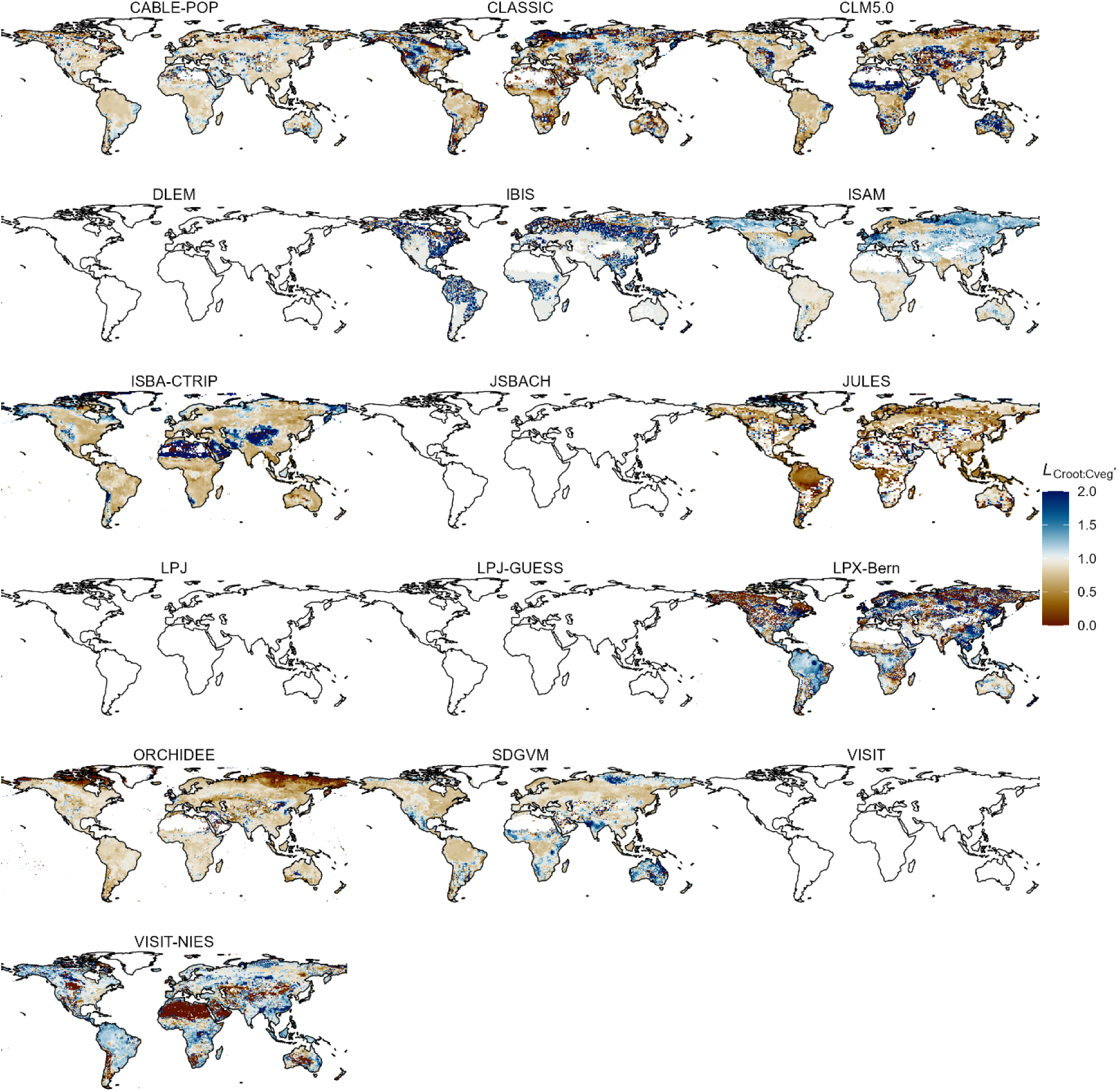
Spatial pattern of *L*_Croot:Cveg*_ for each model. The name of each model is printed on top of each map. *L*_Croot:Cveg*_ = *R*_Croot_ / *R*_Cveg*_ where *R*_Croot_ (*R*_Cveg*_) is the relative change in root C (steady-state total vegetation C), evaluated from simulations with rising CO_2_ and changing N deposition. Both *L* and *R* terms are unitless. As many models include coarse roots in *C*_root_, *C*_root_ should not have rapid response to carbon fertilization. Ideally, we should apply steady state conversion to *C*_root_, and calculate *L*_Croot*:Cveg*_ but this could not be implemented due to the lack of *C*_root_ turnover time. We believe that *L*_Croot:Cveg*_ is less useful than *L*_Croot:Cveg_ (Figure S8).

## 5. Linearity *L*_Cleaf:Cveg*_

**Figure S13.**
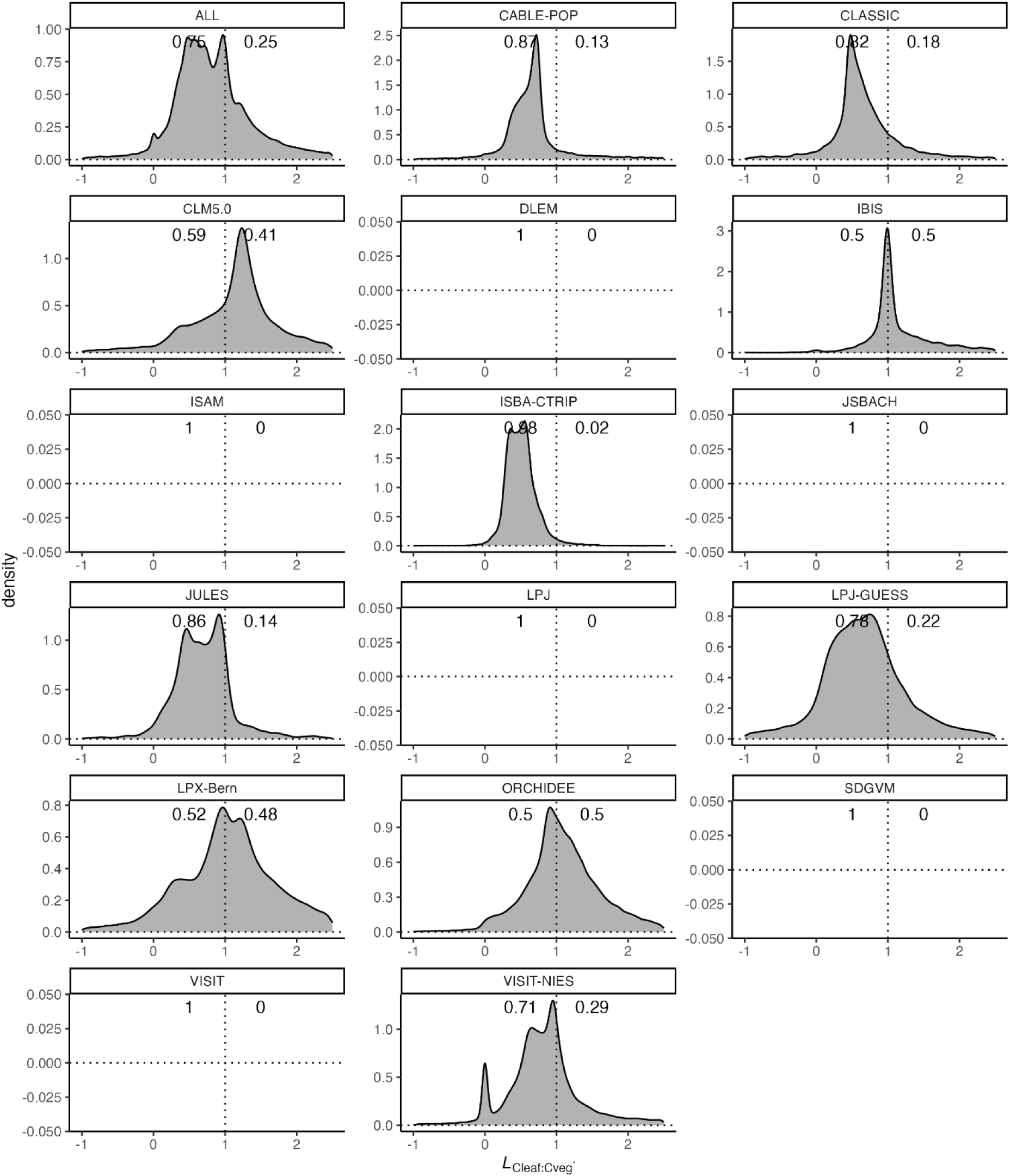
Density of the distribution of *L*_Cleaf:Cveg*_ values across gridcells by model, printed as labels on top of each panel. The proportion of gridcells with *L*_Cleaf:Cveg*_ < 1 (*L*_Cleaf:Cveg*_ > 1) is given by the annotation on the left (right) side of the plotting area. The panel labelled ‘ALL’ represents the joint pattern with data pooled from all models. *L*_Cleaf:Cveg*_ = *R*_Cleaf_ / *R*_Cveg*_ where *R*_Cleaf_ are the relative change of leaf C (steady-state total vegetation C), evaluated from simulations with rising CO_2_ and changing N deposition.

**Figure S14.**
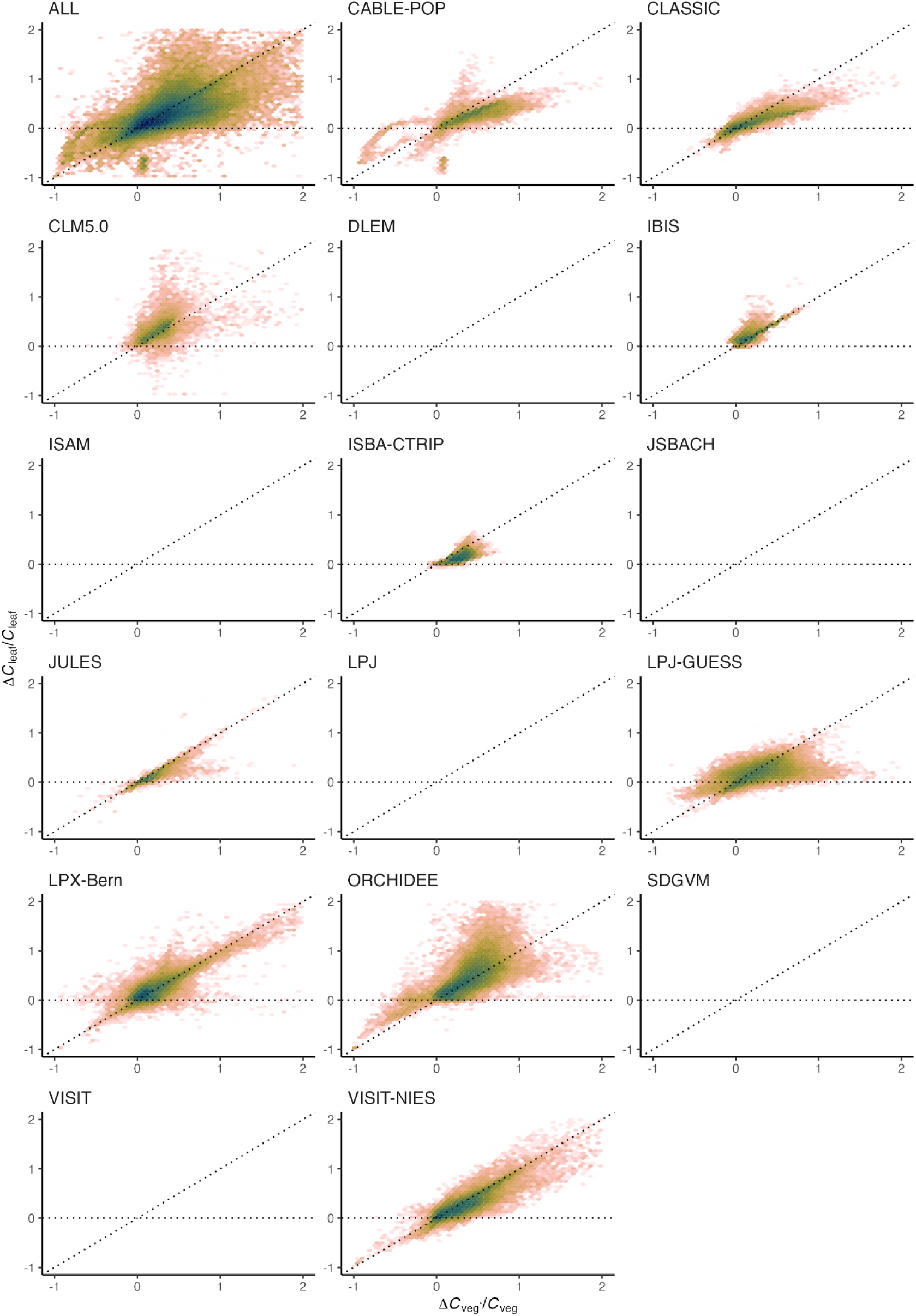
Density scatter plot of *R*_Cleaf_ vs. *R*_Cveg*_ values across all gridcells for each model. The name of the model is printed on top of each histogram. Dark colors denote high density, while bright colors denote low density. *R*_Cleaf_ is the relative change (ΔC_leaf_/C_leaf_, unitless) of leaf C. *R*_Cveg*_ is the relative change (ΔC_veg*_/C_veg_, unitless) of steady-state total vegetation C. Both relative changes are evaluated from the same simulations. The panel labelled ‘ALL’ represents the joint pattern with data pooled from all models.

**Figure S15.**
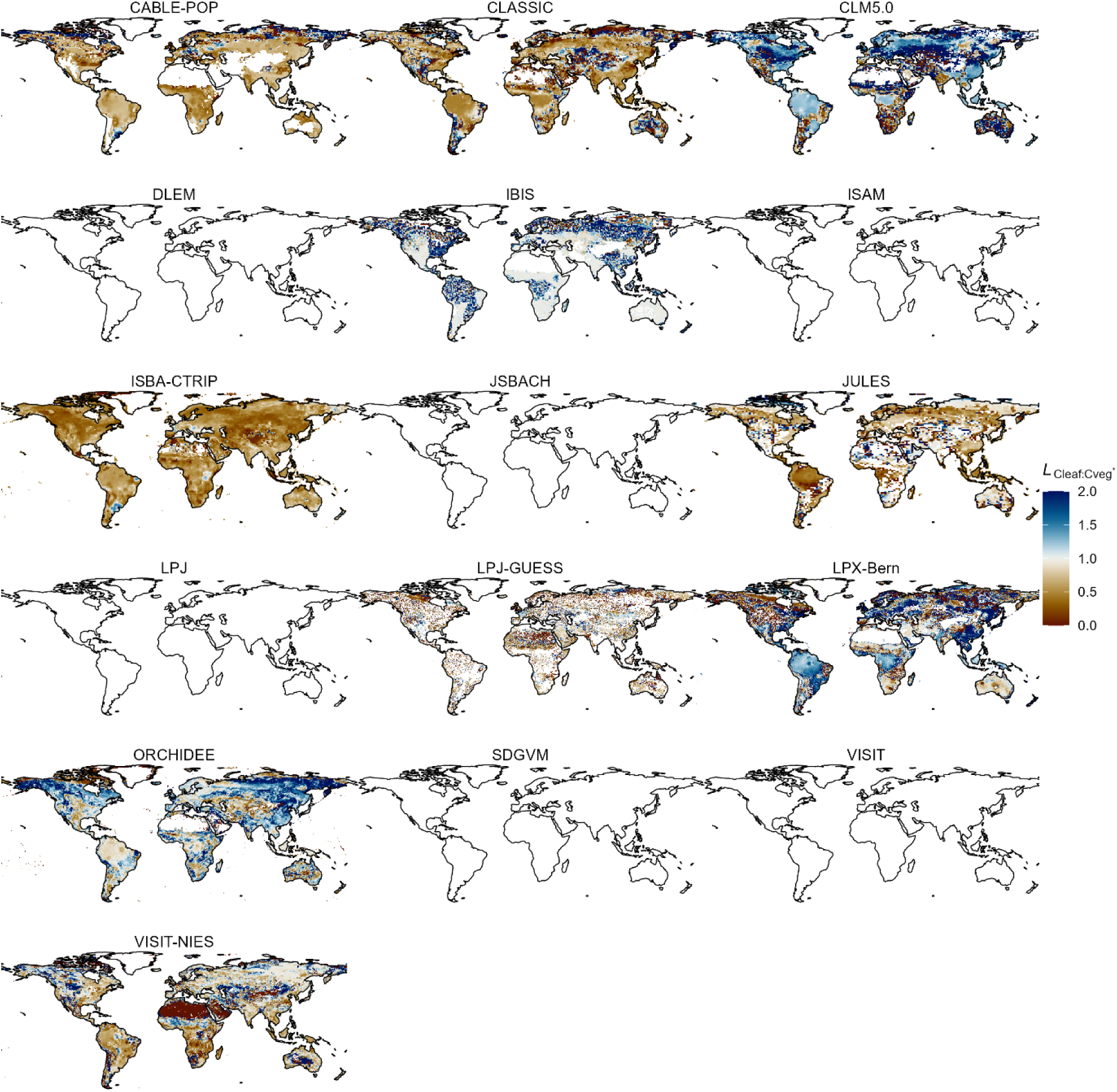
Spatial pattern of *L*_Cleaf:Cveg*_ for each model. The name of each model is printed on top of each map. *L*_Cleaf:Cveg*_ = *R*_Cleaf_ / *R*_Cveg*_ where *R*_Cleaf_ (*R*_Cveg*_) is the relative change in leaf C (steady-state total vegetation C), evaluated from simulations with rising CO_2_ and changing N deposition. Both *L* and *R* terms are unitless.

## 6. Linearity *L*_Cwood:Cveg_

**Figure S16.**
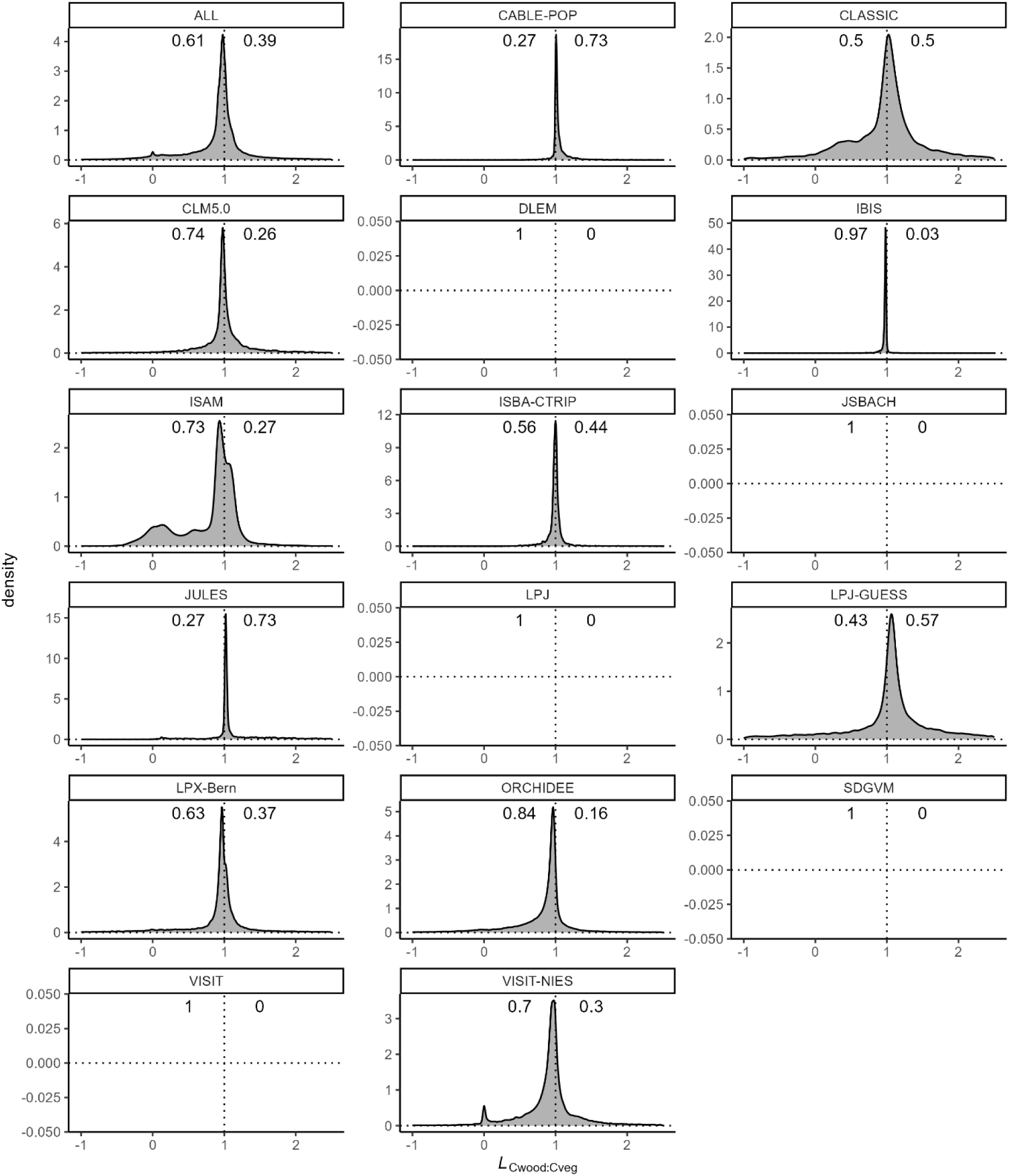
Density of the distribution of *L*_Cwood:Cveg_ values across gridcells by model, printed as labels on top of each panel. The proportion of gridcells with *L*_Cwood:Cveg_ < 1 (*L*_Cwood:Cveg_ > 1) is given by the annotation on the left (right) side of the plotting area. The panel labelled ‘ALL’ represents the joint pattern with data pooled from all models. *L*_Cwood:Cveg_ = *R*_Cwood_ / *R*_Cveg_ where *R*_Cwood_ (*R*_Cveg_) are the relative change of wood C (total vegetation C), evaluated from simulations with rising CO_2_ and changing N deposition. Both *L* and *R* terms are unitless.

**Figure S17.**
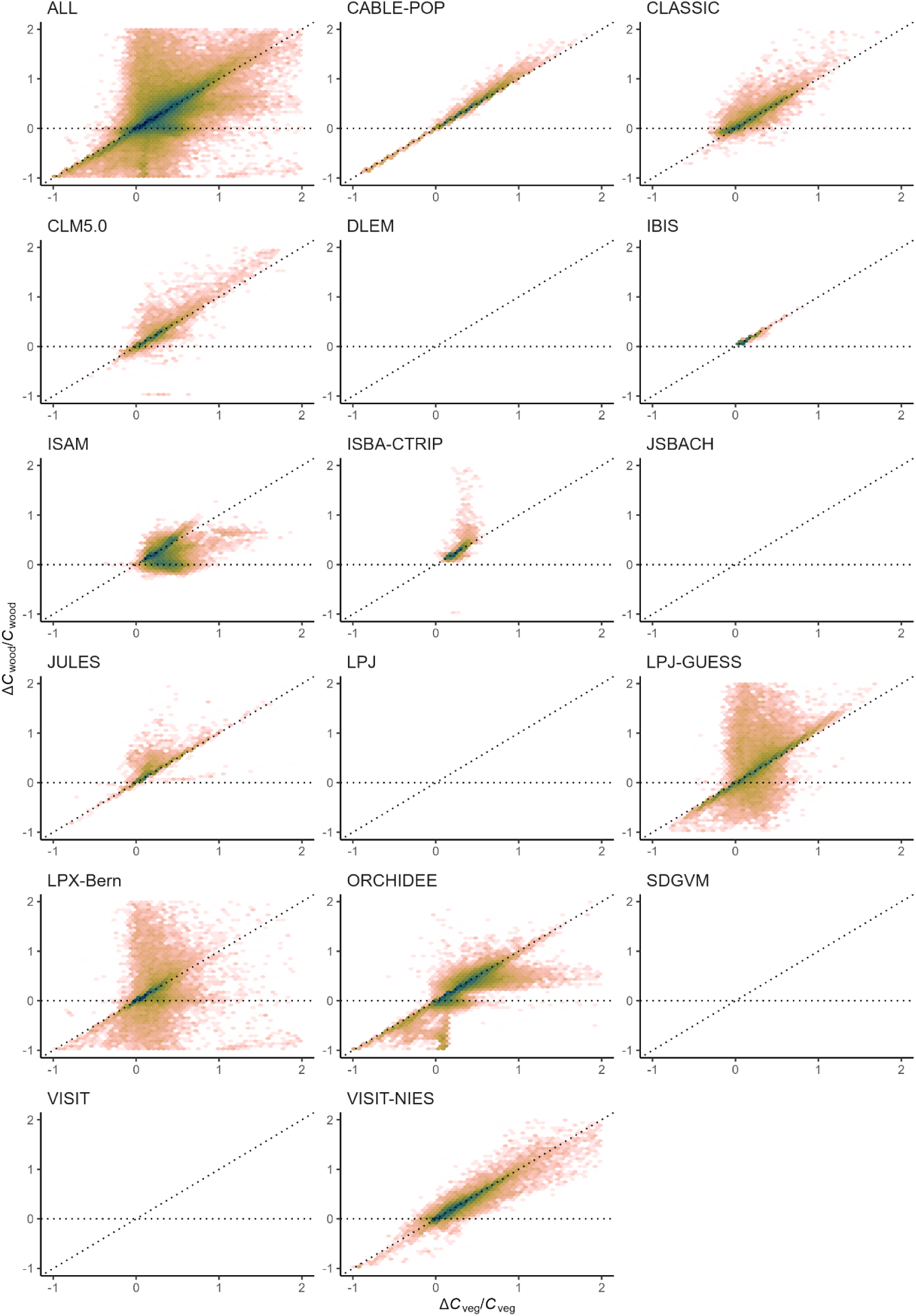
Density scatter plot of *R*_Cwood_ vs. *R*_Cveg_ values across all gridcells for each model. The name of the model is printed on top of each histogram. Dark colors denote high density, while bright colors denote low density. *R*_Cwood_ is the relative change (ΔC_wood_/C_wood_, unitless) of wood C. *R*_Cveg_ is the relative change (ΔC_veg_/C_veg_, unitless) of total vegetation C. Both relative changes are evaluated from the same simulations. The panel labelled ‘ALL’ represents the joint pattern with data pooled from all models.

**Figure S18.**
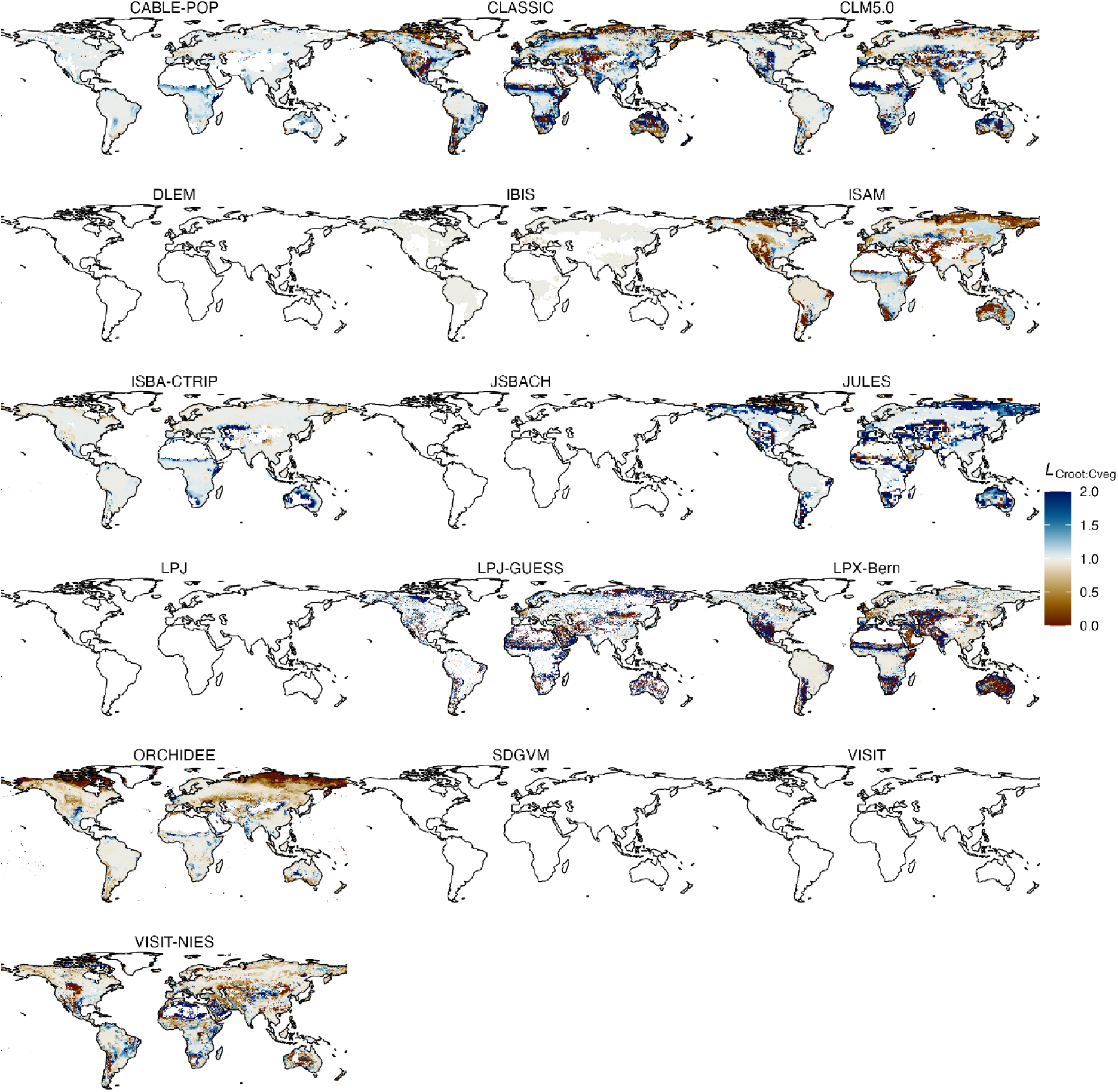
Spatial pattern of *L*_Cwood:Cveg_ for each model. The name of each model is printed on top of each map. *L*_Cwood:Cveg_ = *R*_Cwood_ / *R*_Cve*_ where *R*_Cwood_ (*R*_Cveg_) is the relative change in wood C (steady-state total vegetation C), evaluated from simulations with rising CO_2_ and changing N deposition. Both *L* and *R* terms are unitless.

## 6. Other figures

**Figure S19.**
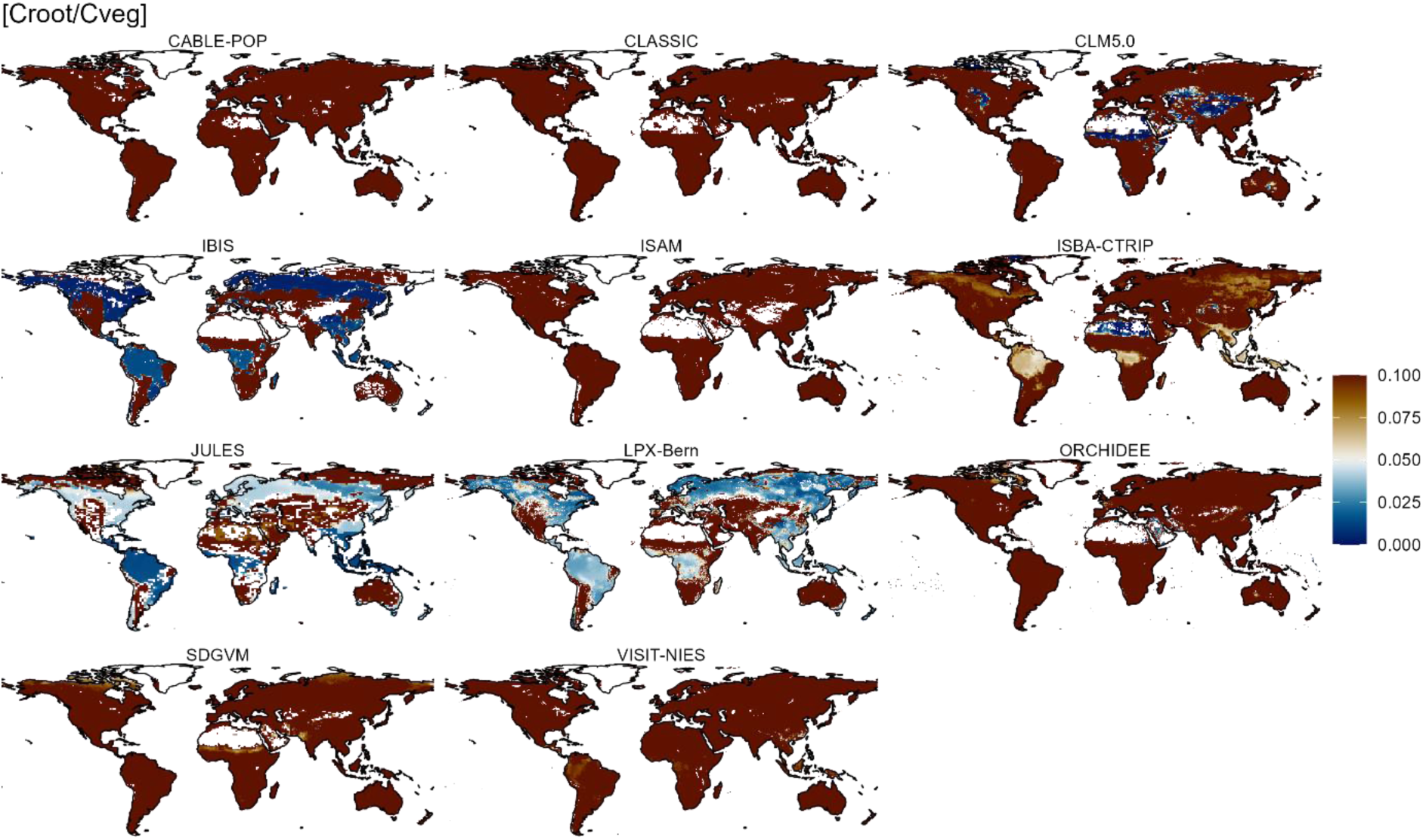
Root biomass (C_root_) divided by vegetation biomass (C_veg_) for each model. This figure is used to diagnose which model includes coarse roots in output variable C_root_. The idea is that for tropical evergreen forests, C_root_ / C_veg_ > 5% would signal that coarse root is included in C_root_. We did try to find out whether the model outputs of C_root_ include coarse roots, by reading manuals and contracting modelers. However, some modelers just can’t remember or they are not responsible for roots simulation. Such information may be archived somewhere in the output explanation of the model manual. In this case, the only risk is that the model manual was not written for TRENDY and the TRENDY outputs could be in a different format. If unfortunately not explained in the output part, it would be somewhere in the codes or equations which will take a whole day to find out (an example of model opaqueness). In any case, the most reliable and efficient way to find out, is by diagnosing the output and draw C_root_ / C_veg_

## Supplementary information for

**Figure S20.**
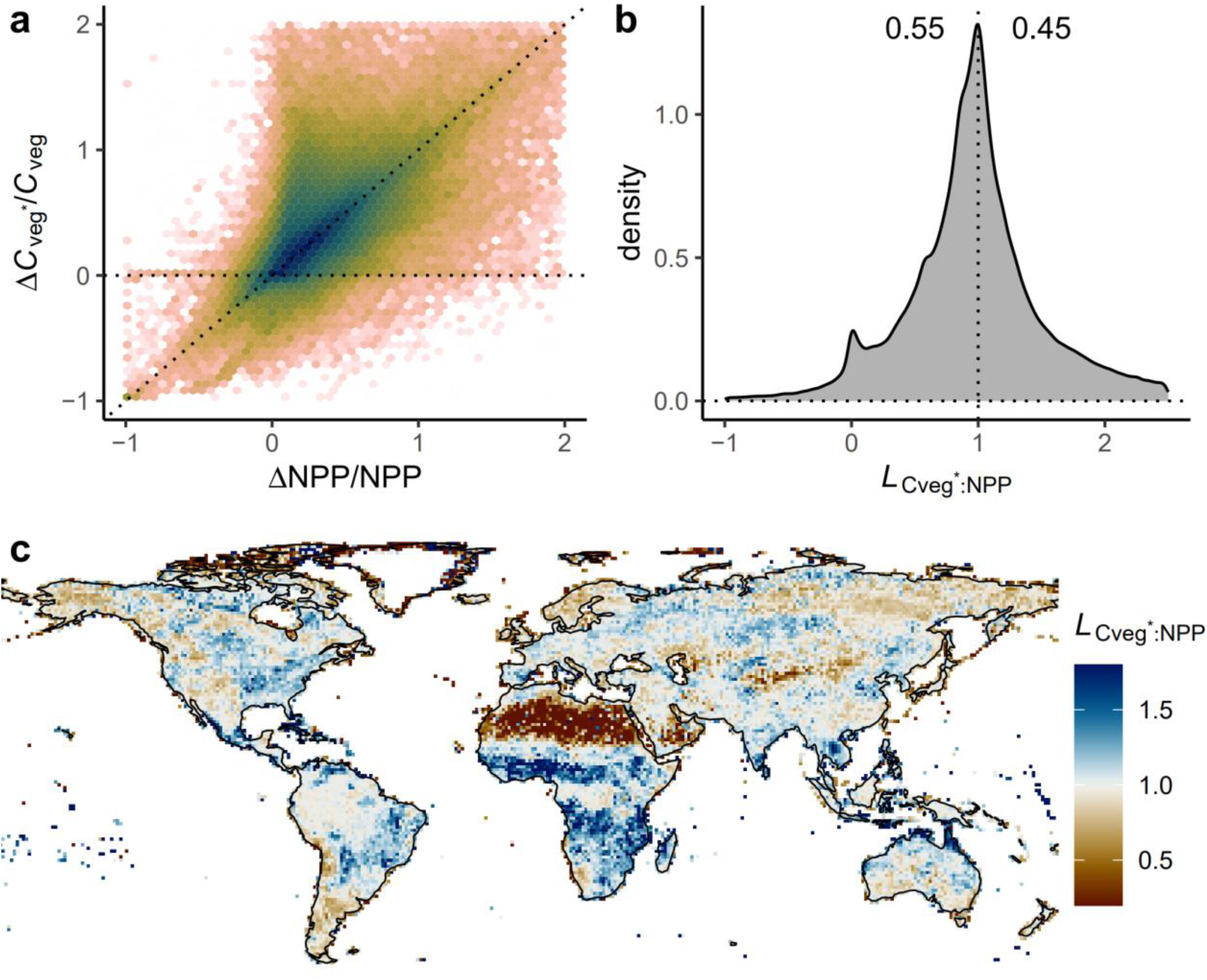
The relationship between the relative change of steady-state vegetation C (*R*_Cveg*_ = ΔC_veg*_/C_veg_) and the relative change of net primary productivity (*R*_NPP_ = ΔNPP/NPP). Same figure as Figure 3 (in the main text) but here we include gridcells with C_veg_ lower than 5% quantile (i.e. include sparse desert). By comparing with Figure 3, it could be concluded that small C_veg_ values lead to a spike at of *L*_Cveg*:NPP_ = 0.

**Supplementary Data 1.**
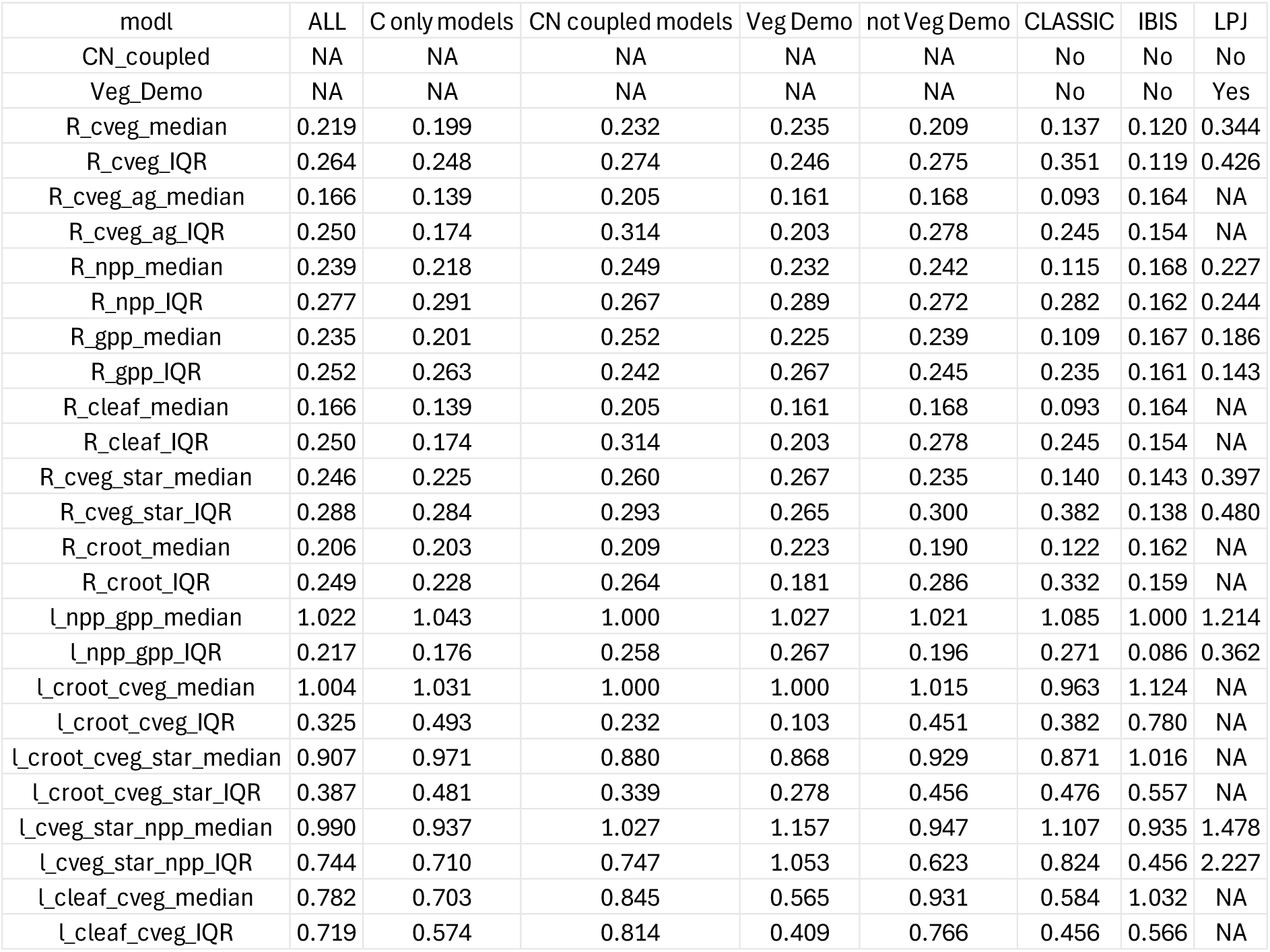
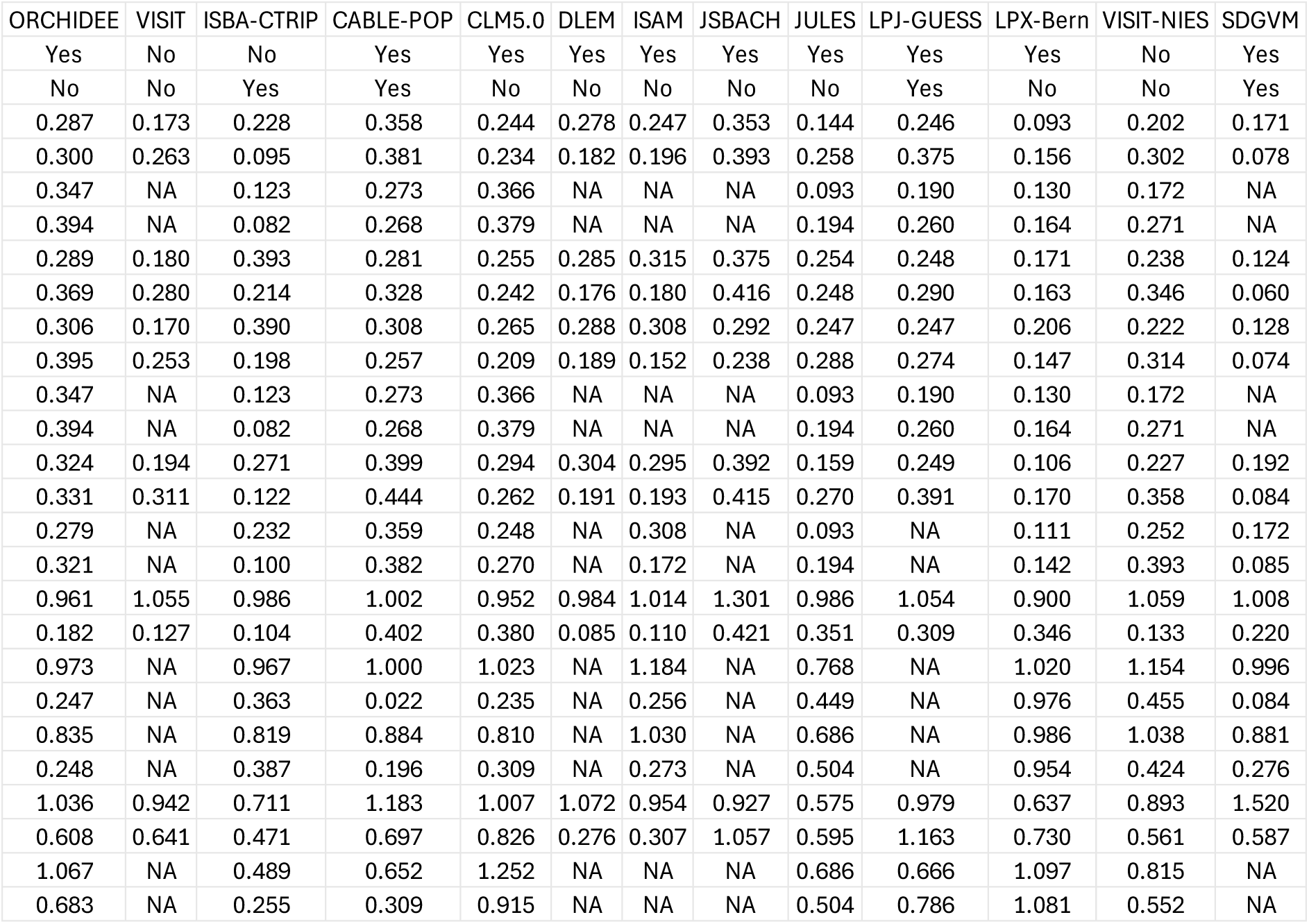
for peer-review. Median and inter-quantile range (IQR) for each relative change and linearity term. This table is inserted here as a screenshot for peer-reviewing purpose. This table will be removed upon acceptance, then the table will be available as Supplementary Data.

